# Human Th17-state classification reveals a selective vulnerability in acylcarnitine metabolism in type 2 diabetes

**DOI:** 10.64898/2026.09.01.748572

**Authors:** Naveena S. Ujagar, Suresh Poudel, Shubh Saraswat, Suhas Sureshchandra, Josh Kim, Uyen-Vy Le, Albert R. Jones, Nicole Hopkins, Trupti Sriram, Matthew Matson, Emely H. Pilier, Marlyd Mejia, Samuel Bailin, Celestine Wanjalla, Micheal Sy, Dawn Newcomb, Craig Walsh, Lisa E. Wagar, Barbara S. Nikolajczyk, Xiaohua D. Zhang, Doug Green, Dequina Nicholas

## Abstract

Th17 cells are categorized as non-pathogenic (Th17np) and pathogenic (Th17p), states defined primarily in murine autoimmune models. In contrast, studying these states in humans remains difficult because these subsets lack validated identifiers. We developed a nanovial-based single-cell ELISA to isolate cytokine-secreting human Th17 cells and define Th17np vs Th17p cell transcriptional states. These signatures enabled predictive classification and flow-cytometric identification across tissues and diseases. Application of these strategies with single-cell metabolic modeling and functional profiling revealed that Th17np cells predominate in type 2 diabetes yet exhibit impaired fatty acid oxidation capacity despite elevated carnitine palmitoyltransferase 1A (CPT1A). CRISPR perturbation and pharmacologic inhibition identified acylcarnitine metabolism as a regulator of Th17 state balance and exposed a selective metabolic vulnerability in type 2 diabetes. This framework links human Th17 cell pathogenicity to lipid metabolism.

## Introduction

Th17 cells maintain mucosal barriers and protect against extracellular bacterial and fungal infections but can also drive inflammatory and autoimmune disease^1–7^. Much of this functional heterogeneity has been resolved using *in vitro* differentiation and murine autoimmune models to classify non-pathogenic (Th17np) and pathogenic (Th17p) cell states. The transcription factor retinoid-related orphan receptor gamma-t (RORγt), is the master transcriptional regulator of Th17 cells resulting in production of IL-17A, IL-17F, and IL-21^8–12^.Th17 differentiation is regulated by cytokines including TGFβ, IL-6, and IL-21, with IL-23 critical for pathogenicity^13–17^. In humans, however, Th17np and Th17p cell states cannot be directly identified and functionally compared because primary human Th17 subsets lack validated markers capable of distinguishing these populations.

Murine models of autoimmunity are foundational to our current understanding of Th17 cells and their pathogenicity; however, much of that knowledge has not been directly translated to humans. Pioneering studies demonstrated that Th17 cells are not a fixed lineage but exhibit significant plasticity, converting to a Th1-like phenotype with co-expression of IFN-γ in inflammatory environments *in vivo* raising the question of whether functional heterogeneity existed within the Th17 compartment itself^18–20^. Transcriptional profiling of *in vitro* differentiated cells and CNS-infiltrating Th17 cells during EAE identified gene programs associated with each state and subsequent studies demonstrated that regulators including CD5L and TCF1 can restrain differentiation to a Th17p state^21–23^. Despite these mechanistic advances, no validated molecular or phenotypic framework directly resolves Th17np and Th17p states in primary human samples. Human Th17 pathogenicity is therefore generally inferred from the disease or inflammatory context in which the cells are found rather than measured as an intrinsic cell state.

Cellular metabolism is an established mechanism governing Th17 differentiation and pathogenicity^24–26^. The differentiation of naïve T cells into Th17np vs Th17p is dependent on ACC1, while Gpi1 is selectively required for Th17p cell in hypoxic environments, creating the possibility of Th17 state selective intervention. One clear example is the discovery of polyamine metabolism regulating murine Th17 differentiation to a pathogenic state, highlighting the polyamine pathway as a potential target for autoimmune therapies. Discovery of this state-specific metabolic program was enabled by a metabolic flux algorithm based on single-cell transcriptomic data called COMPASS^27^. Together, these first studies to successfully target Th17p through metabolic enzymes to reduce their pathogenicity were discoveries made primarily through *in vitro* polarization and murine models. Therefore, their relevance to primary human Th17 states is unknown. Determining whether metabolic dependencies govern human Th17 pathogenicity requires both identification of the pathogenic states and recovery of sufficient primary cells for transcriptomic and functional analysis.

Several challenges exist to translating our understanding of the metabolic phenotypes of Th17 cells described *in vitro* and in murine models to human, thus precluding targeting Th17p versus Th17np cells in human disease. Th17 cells comprise only 0.5 to 2% of CD4+ T cells and only 0.1 to 1% of circulating peripheral blood mononuclear cells (PBMCs)^28,29^. Heterogeneous cultures dilute Th17-specific responses, and cytokine-driven skewing of naive CD4+ T cells may not faithfully replicate the transcriptional and metabolic phenotypes shaped by the *in vivo* environment. Most critically, existing approaches cannot directly and reliably assign pathogenic state to primary human Th17 cells for matched transcriptional, metabolic and functional analysis.

Here, we established complementary single-cell transcriptomic and flow-cytometric frameworks for identifying human Th17 pathogenicity states. A nanovial-based single-cell ELISA enabled enrichment and transcriptional definition of cytokine-secreting Th17 cells from primary human samples, generating signatures that classified Th17 states across tissues and diseases. Integration of these frameworks with metabolic modeling and functional profiling revealed that Th17np cells predominate in type 2 diabetes despite impaired fatty acid oxidation capacity and elevated CPT1A. CRISPR perturbation and pharmacologic inhibition further identified acylcarnitine metabolism as a regulator of Th17-state balance and a selective metabolic vulnerability in type 2 diabetes.

## Results

### Single cell ELISAs enrich Th17 cells by cytokine secretion and allow identification of Th17 cell heterogeneity

To facilitate the study of viable human Th17np and Th17p cells, we developed a novel Th17 enrichment assay to capture primary human Th17 cells **(Fig. 1A)**. Hydrogel nanovials were functionalized by non-covalently binding antibodies to the cell capture surface via a biotin-streptavidin interaction. We used anti-human CD45 to capture pre-enriched and anti-CD3/CD28 activated CD4+ T cells and used anti-human IL-17A, IL-17F, and IL-21 to locally capture these Th17-secreted cytokines **(Fig. 1B)**. Use of dual magnetic and fluorescent labeled secondary antibodies against IL-17A, IL-17F and IL-21 effectively create single cell ELISAs that allow for magnetic isolation of Th17 cells. We confirmed demonstrate that Th17 cells secreting at least one of the three cytokines were viable and enriched to 90.93% of cells **(Fig 1C-D, Supplemental Fig. 1A-D**).

**Figure 1.**
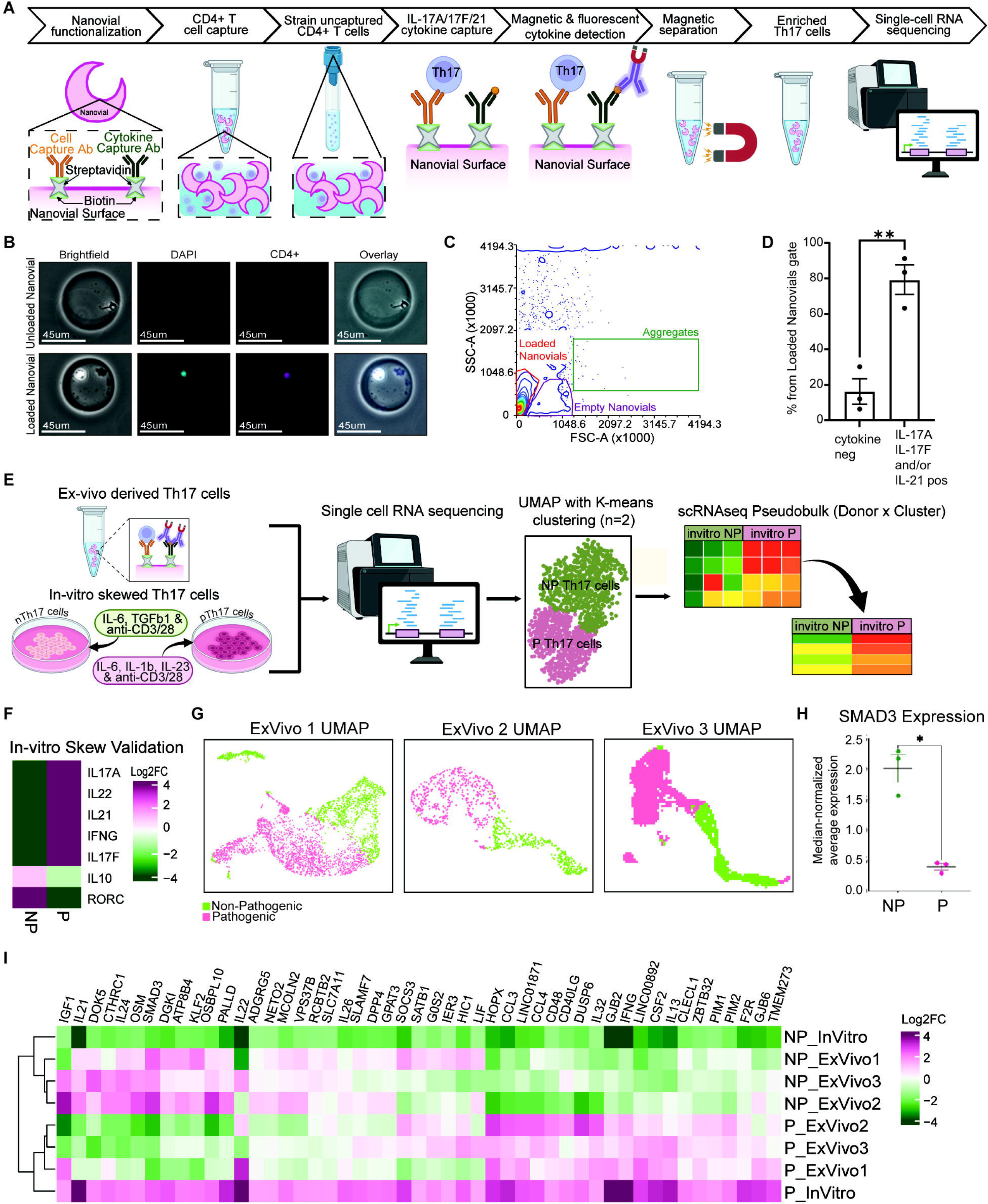
Nanovial-based single-cell ELISA enriches viable cytokine-secreting human Th17 cells for transcriptomic analysis. (**A**) Experimental workflow for nanovial enrichment of Th17 enrichment and scRNA-seq. (**B**) Representative bright-field and fluorescence images of unloaded and CD4+ T-cell-loaded nanovials. (**C**) Representative flow cytometry discrimination of empty, cell-loaded, and aggregated nanovials. (**D**) Percentage of loaded nanovials negative or positive for at least one Th17-associated cytokine (IL-17A, IL-17F, or IL-21; n = 3 donors). (**E**) Experimental workflow for matched scRNA-seq of nanovial-enriched *ex vivo* Th17 cells and *in vitro*-skewed Th17np and Th17p cells, followed by k-means clustering and pseudobulk analysis. (**F**) Heatmap of canonical cytokine- and lineage-associated genes in *in vitro*-skewed Th17np and Th17p cells. (**G**) UMAP plots showing k-means classification of nanovial-enriched Th17np (green) and Th17p (magenta) cells from three independent donors. (**H**) Median normalized average SMAD3 expression in nanovial-enriched Th17np and Th17p cells from three donors. (**I**) Unsupervised hierarchical clustering of pseudobulk gene expression in *in vitro*-skewed and nanovial-enriched Th17np and Th17p cells. In (D) and (H), bars or horizontal lines show mean ± SEM and points represent donors. Statistical significance was determined by two-tailed t test; *P < 0.05 and **P < 0.01.

Once we confirmed the ability to enrich viable Th17 cells, we asked whether transcriptomic signatures of Th17 cells skewed *in vitro* were the same as Th17 cells derived *ex vivo*. From three lean healthy donors, naïve CD4+ T cells were skewed *in vitro* to Th17np and Th17p cells and endogenous Th17 cells were enriched using our nanovial scELISA method. We then performed single-cell RNA sequencing (scRNA-seq) on these matched Th17 cell samples **(Fig. 1E)**. *In vitro* skewed cells exhibited expected difference in key genes with Th17p cells expressing higher transcripts of IL17 and Th17np expressing more RORC **(Fig. 1F)**. Nanovial-enriched Th17 cells were classified into two subpopulations, Th17np and Th17p, using K-means clustering (k=2). A key distinguishing feature of this classification was the elevated expression of SMAD3 in Th17np cells relative to Th17p cells, consistent with previously reported *in vitro* findings^30,31^ **(Fig. 1G-H)**. Pseudobulk was performed on the *in vitro* skewed Th17np and Th17p samples to reduce biological variability inherently introduced by non-skewed cells^32^. A heatmap of the top 50 differentially expressed genes from the *in vitro* differentiated Th17p samples were produced after k-means clustering of the nanovial enriched samples and pseudobulking of *in vitro* skewed samples **(Fig 1I, Supplemental Fig. 1E)**. Unbiased hierarchical clustering analysis of the samples revealed that *in vitro* Th17np and Th17p groups clustered with their respective endogenous counterparts **(Fig. 1I)**. The unbiased clustering of these data demonstrated that Th17np and Th17p cells cluster separately as expected, and can be defined as Th17p cells based on published^33–35^ genes for these cell type in mice including IFNG, CSF2, CCL4, and IL22 **(Suppl 2F, Fig 1H)**. However, these data also highlight stark differences of *in vivo* differentiated Th17 cells that fail to be recapitulated *in vitro,* suggesting that metabolic differences between the source of Th17 cells may also exist. We conclude that the scELISA nanovial enrichment method can capture viable, Th17 cells actively secreting IL-17A/F/21+, thereby enabling analysis of the human Th17np and Th17p transcriptome.

### A single cell framework identifies classifies Th17np & Th17p cells by their transcriptome and through flow cytometry

From these data, we devised two approaches to identify primary human Th17 cell states. First, we developed a predictive model that could annotate human Th17 cells as Th17np or Th17p. The model was trained on scRNA-sequencing data from *in vitro* differentiated Th17 cells. scRNA-seq data from *in vitro* skewed and nanovial-enriched samples underwent quality control to remove ambient RNA, doublets, and low-quality cells. Then scVI was used for batch correction and to harmonize the data across technical batches, and subsetted for CD4+CD161+ cells to exclude naïve cells from the *in vitro* skewed cells and ensure Th17 lineage^36^ **(Fig. 1A)**. For classification, we applied SCANVI^37^, a semi-supervised neural network that trained on 8,000 of the most highly variable genes in the *in vitro* skewed cells. To ensure unbiased performance estimates, we implemented 5-fold cross-validation. This approach achieved ∼94% accuracy with only ∼2% ambiguous cells **(Supplemental Fig. 2A-B)**. The decision boundary for nanovial-enriched cells showed clear separation with most cells exhibiting high confidence in either Th17p or Th17np classification **(Fig 2B)**. When applied to nanovial-enriched cells, the model produced distinct clusters of pathogenic, non-pathogenic, and a small population of ambiguous cells **(Fig 2C)**.

**Figure 2.**
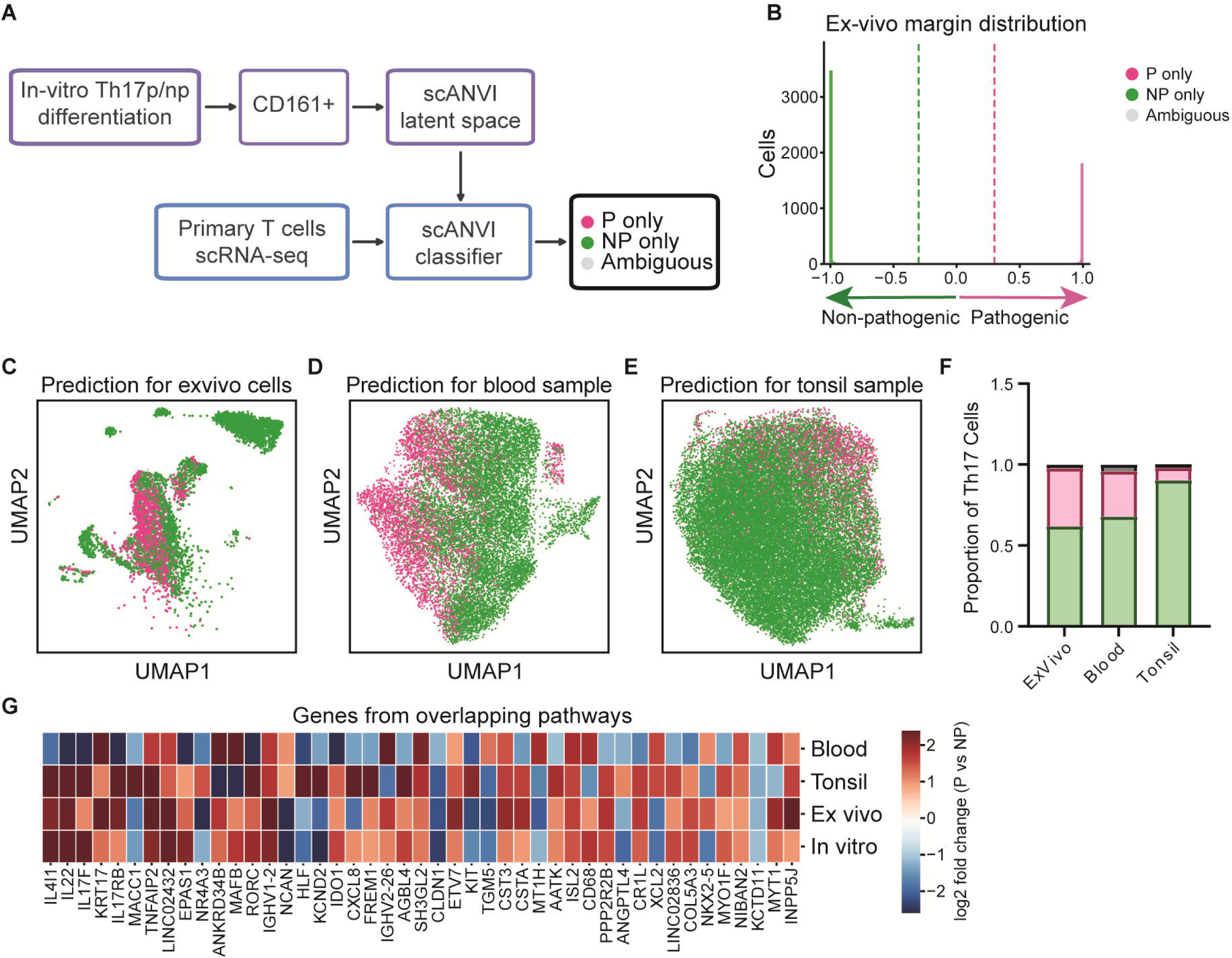
A transcriptomic classifier identifies human Th17np and Th17p states across tissues. (**A**) Schematic of the scANVI classification framework. The model was trained in the scVI latent space using labeled CD161+ *in vitro*-skewed Th17np and Th17p cells and applied to primary T cell scRNA-seq datasets to classify cells as Th17np, Th17p, or ambiguous. (**B**) Classifier-margin distribution for nanovial-enriched *ex vivo* Th17 cells. Dashed lines indicate the decision thresholds for Th17np only (green), Th17p only (magenta), and ambiguous (gray) classifications. (**C to E**) UMAP plots showing predicted Th17np, Th17p, and ambiguous cells in nanovial-enriched *ex vivo* samples (C), peripheral blood (D), and tonsil (E). (**F**) Proportions of classified Th17np, Th17p, and ambiguous cells in *ex vivo*, blood, and tonsil datasets. (**G**) Heatmap showing the log2 fold change in expression between Th17p and Th17np cells for genes associated with pathways shared across blood, tonsil, *ex vivo*, and *in vitro* datasets.

We validated model robustness using large, independent blood and tonsil transcriptomic datasets^38^. After subsetting for CD4⁺CD161⁺ cells, we adapted the trained model to new data using scArches^39^, which efficiently fine-tunes the model without retraining from scratch. The classifier performed consistently across tissues. Blood samples yielded 10,923 Th17np cells, 4,469 Th17p cells, and 4.2% ambiguous. Tonsil samples yielded 29,188 Th17np cells, 2,513 Th17p cells and 2.0% ambiguous **(Fig 2D-E)**. This consistent performance across independent donors and tissue types demonstrates the model’s generalizability.

To identify molecular features distinguishing Th17np from Th17p states, we performed differential expression analysis using stringent criteria (Bayes factor ≥3, |log₂ fold-change| ≥1). We observed substantial overlap of differentially expressed genes across all four datasets (*in vitro*, *ex vivo*, blood, and tonsil) with key Th17-associated genes (*IL22, IL17F,* RORC) showing consistent directional changes, with the exception of sequenced blood which differs from our anti-humanCD3/28 stimulated *ex vivo* samples possibly due to lack of stimulation **(Fig 2G)**. Notably, several canonical Th17 transcription factors (*MAFB*, *NR4A3*) were differentially expressed, along with inflammatory mediators (*CXCL8*, *TNFAIP2*) and tissue-remodeling factors (*KRT17*, *TGM5*, *CLDN1*). Lower gene expression in Th17p cells included *LINC0109*, *CLDN*, *KCTD11*, and *AC005208.1* **(Supplemental Fig. 2C)**. Conversely, 9 genes had higher expression in pTh17 cells, including transcription factors *ETV7* and *ISL2*; which have not been previously described in CD4+ T cell literature **(Supplemental Fig. 2D)**. We conclude that the reproducibility of these transcriptomic signatures across diverse biological sources suggests that Th17np and Th17p cells maintain core molecular programs regardless of tissue environment, and that this predictive model reliable annotates Th17np vs Th17p cells via transcriptomic signatures.

We next leveraged the Th17 scSeq data to develop a flow cytometry gating strategy to allow for scalable identification of Th17 states. Differential gene expression analysis of *in vitro* Th17np and Th17p samples identified *CD101* and *TIMD4* as statistically significantly differentially expressed surface marker-encoding genes, however, flow cytometry of primary lean donor samples revealed few detectable CD101+ and TIM4+ events with notably limited IL-17A/F secretion relative to the broader CD3+CD4+ T cell population **(Supplemental Fig. 3)**, prompting exploration of alternative markers. Therefore, we used markers of functionality to discriminate between Th17np and Th17p cells. The hallmark IFN-γ response was strongly upregulated in *in vitro* skewed Th17p cells (NES = 2.2, FDR < 0) **(Fig 3A)**. We validated these transcriptional findings by skewing naïve CD4+ T cells from lean donors (n = 4) to Th17p and Th17np cells and performing flow cytometry. We found that there was a statistically significant increase of IFN-γ+ CD4+ T cells in the Th17p versus Th17np cells **(Fig 3B)**. This was not due to differences in the proportions of CD4+ T cells in Th17np and Th17p cell samples **(Supplemental Fig. 4A)**. We checked for other cytokines traditionally associated with Th17 cells and found that only IL-17F, not GM-CSF, IL-21 or IL-17A, was also increased in CD4+ T cells skewed toward Th17p versus Th17np states **(Fig 3C-F)**. Therefore, we deemed IFN-γ+ IL-17F+ cells to be sufficient for identifying Th17p cells as these were also increased in CD4+ T cells **(Fig 3G)**. In contrast, we proposed IFN-γ-IL17A+ CD4+ T cells as Th17np cells as there was a trending increase of IL-17A in Th17np cells at a sizable percentage, as compared to IFN-γ+ GMCSF+ or IL-21+ IL-22+ cells **(Suppl 4B-C)**.

**Figure 3.**
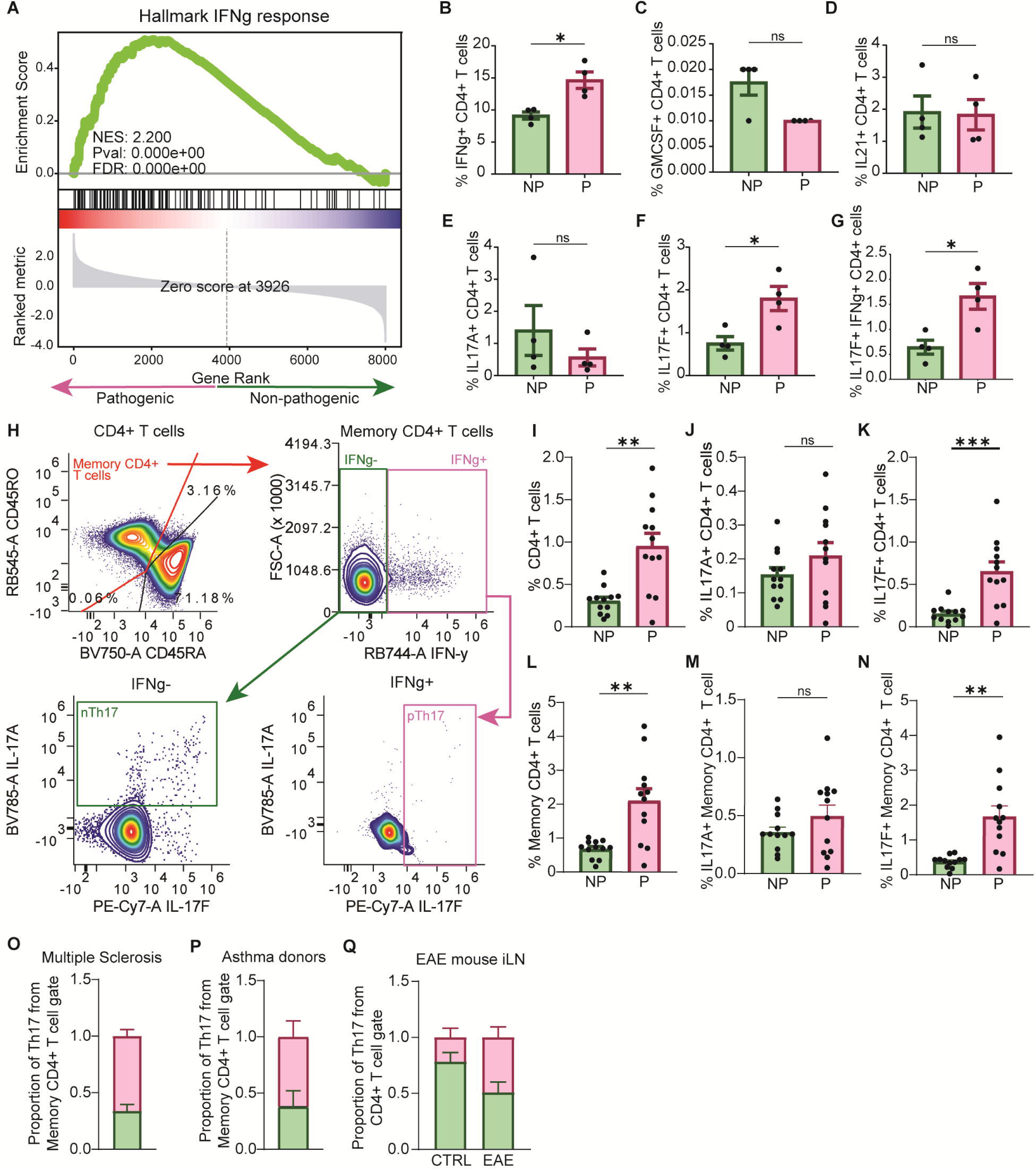
Cytokine-based flow cytometry distinguishes human Th17np and Th17p cells. (**A**) Gene set enrichment analysis of the Hallmark IFN-γ-response pathway in *in vitro*-skewed Th17p relative to Th17np cells. NES, normalized enrichment score; FDR, false-discovery rate. (**B to G**) Frequencies of IFN-γ+ (B), GM-CSF+ (C), IL-21+ (D), IL-17A+ (E), IL-17F+ (F), and IL-17F+IFN-γ+ (G) cells among CD4+ T cells differentiated under Th17np or Th17p conditions (n = 4 donors). (**H**) Representative gating strategy identifying memory CD4+ T cells and defining Th17np cells as IFN-γ−IL-17A+ and Th17p cells as IFN-γ+IL-17F+. (**I to N**) Frequencies of Th17np and Th17p cells among total CD4+ T cells (I), IL-17A+ total CD4+ T cells (J), IL-17F+ total CD4+ T cells (K), memory CD4+ T cells (L), IL-17A+ memory CD4+ T cells (M), and IL-17F+ memory CD4+ T cells (N) from healthy lean donors. (**O and P**) Proportions of Th17np and Th17p cells among memory CD4+ T cells from donors with multiple sclerosis (O) or allergic asthma (P). (**Q**) Proportions of Th17np and Th17p cells among CD4+ T cells from inguinal lymph nodes of control mice and mice with experimental autoimmune encephalomyelitis (EAE). Bars show mean ± SEM; points represent independent donors or biological replicates. Statistical significance was determined by two-tailed t test; *P < 0.05, **P < 0.01, ***P < 0.001; ns, not significant.

The *in vitro* skewed data suggested that Th17p and Th17np cells can be defined based on cytokine secretion, prompting development of a gating strategy that was applied to CD4+ samples from lean donors (n=12) **(Fig 3H, Suppl 4D)**. Th17p cells represented a higher proportion of CD4+ and memory CD4+ T cells than Th17np cells, with no difference in IL-17A+ cell frequency but a statistically significant increase in IL-17F+ cells in Th17p compared to Th17np populations, consistent with *in vitro* findings **(Fig 3I-N)**. To further validate this strategy, we applied it to PBMC samples from patients with established Th17p-driven diseases, multiple sclerosis^3,4^ and allergic asthma^40^, and found a preponderance of Th17p cells **(Fig 3O-P)**; similarly, inguinal lymph node samples from an EAE mouse model, where Th17p inflammation was first described^41,42^, showed a greater proportion of Th17p cells compared to control mice **(Fig 3Q)**, collectively confirming the strategy’s ability to discriminate Th17np from Th17p cells in both human and murine contexts.

### Metabolic phenotyping of Th17np and Th17p cells predicts acylcarnitine metabolism as a metabolic signature discriminating between pathogenic states

With a framework to annotate and identify Th17 pathogenicity both transcriptomically and by flow cytometry, we performed metabolic phenotyping to determine pathways which discriminate between states. First, we applied COMPASS^27^, an algorithm to predict the metabolic state of cells from transcriptomic data at single cell resolution, to our transcriptomic data annotated as Th17np and Th17p. COMPASS revealed that the nucleotide salvage pathway, which implicates N-glycan branching, had strong relationships (Cohen’s d > 0.5) with Th17p cells from all sources **(Fig 4A-B, Supplemental 5A-B)**. This prediction was validated by detection of increased complex N-glycan branching on human Th17p cells compared to Th17np cells **(Supplemental Fig. 6A-B)**. Conversely, COMPASS strongly predicted arachidonic acid metabolism as associated with Th17np cells (Cohen’s d > 0.5) **(Fig 4A-B, Supplemental 5A-B)**. Analysis of core metabolic pathways revealed that fatty acid oxidation had the most striking differences between Th17np and Th17p states **(Fig 4C-D, Suppl 5C-D)**. In particular, multiple enzymes and steps of long chain lipid metabolism, including acyl-CoA ligase and acylcarnitine production and shuttling had a strong relationship (Cohen’s d > 0.5) with Th17np cells from all four sample sources. Collectively, COMPASS analysis revealed that Th17np cells consistently displayed strong signatures of long chain lipid metabolism.

**Figure 4.**
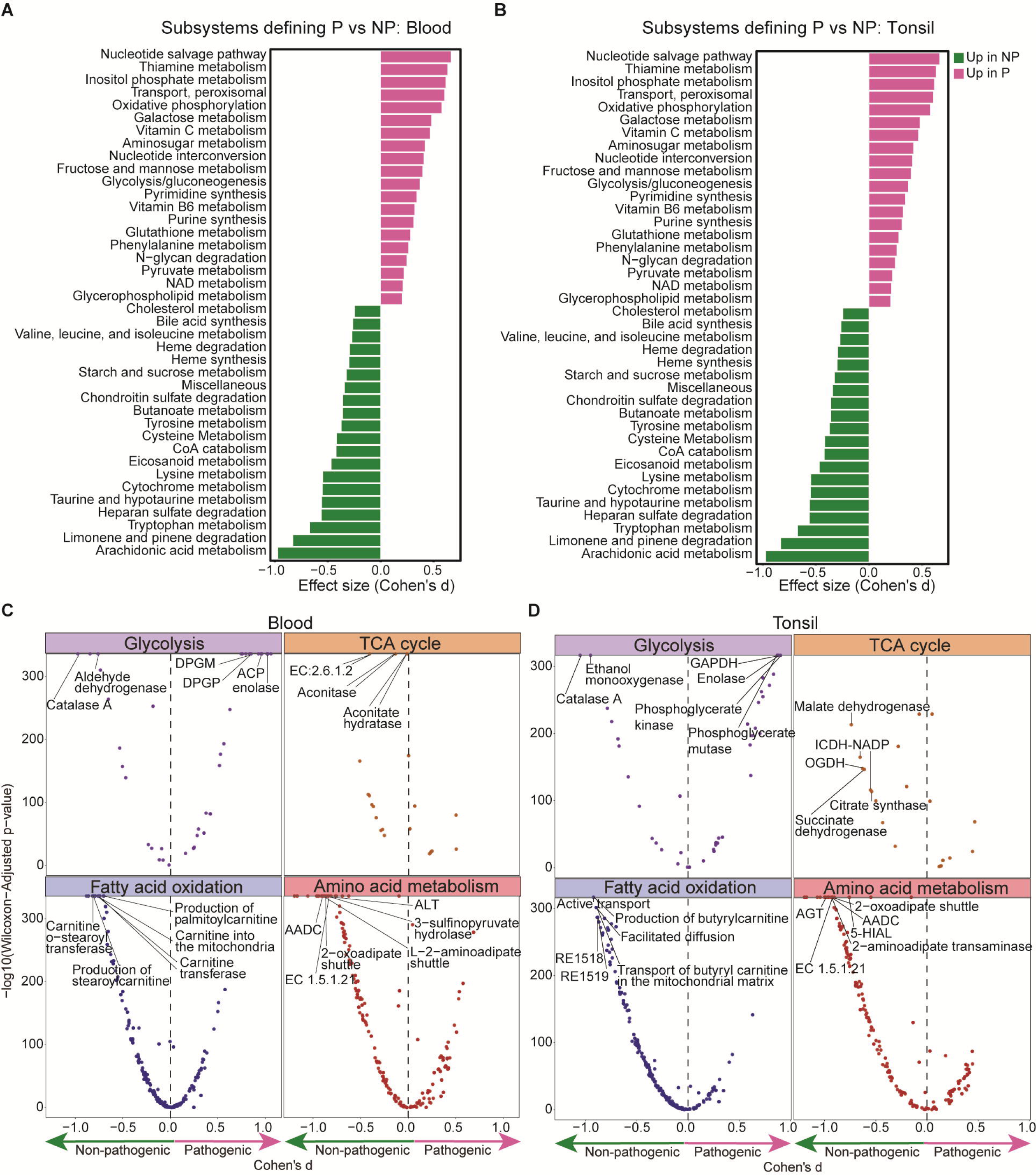
COMPASS predicts distinct metabolic programs in human Th17np and Th17p cells. (**A and B**) COMPASS-predicted metabolic subsystems distinguishing Th17p from Th17np cells in peripheral blood (A) and tonsil (B). Bars show Cohen’s d effect sizes. Positive values (magenta) indicate pathways associated with Th17p cells, and negative values (green) indicate pathways associated with Th17np cells. (**C and D**) Reaction level differences between Th17p and Th17np cells across glycolysis, the tricarboxylic acid (TCA) cycle, fatty acid oxidation, and amino acid metabolism in blood (C) and tonsil (D). Dashed lines indicate effect-size thresholds.

To test the prediction that Th17np cells are associated with long chain lipid metabolism, we applied our novel gating strategy within the disease context of Type 2 Diabetes (T2D). Patients with obesity-associated T2D experience chronic Th17-driven inflammation and exhibit defects in FAO and lipid metabolism pathways that trigger Th17 cytokine secretion^43^. Therefore, we hypothesized that the lipid rich nutrient environments within persons with obesity and T2D would correspond to differences in fatty acid oxidation capacity of these Th17 cells. To test this hypothesis and derive further understanding of the metabolic programs that differ between Th17np and Th17p cells in the context of T2D, we annotated Th17 cells in published adipose-tissue datasets of lean obese and T2D donors^44,45^ (**Fig. 5A-D**). Surprisingly, adipose tissue has an overwhelming proportion of Th17np cells. Further, individuals with T2D had virtually no pTh17 cells compared with lean and obese donors, indicating that T2D-associated Th17 inflammation is dominated by nTh17 cells. We validated this observation with flow cytometry on CD4+ T cells from an independent cohort of normoglycemic lean (BMI≤28), normoglycemic obese (BMI>28), and obese T2D donors combining two published metabolic flow cytometry panels: SCENITH and Met-flow **(Supplemental Table 3)**.

**Figure 5.**
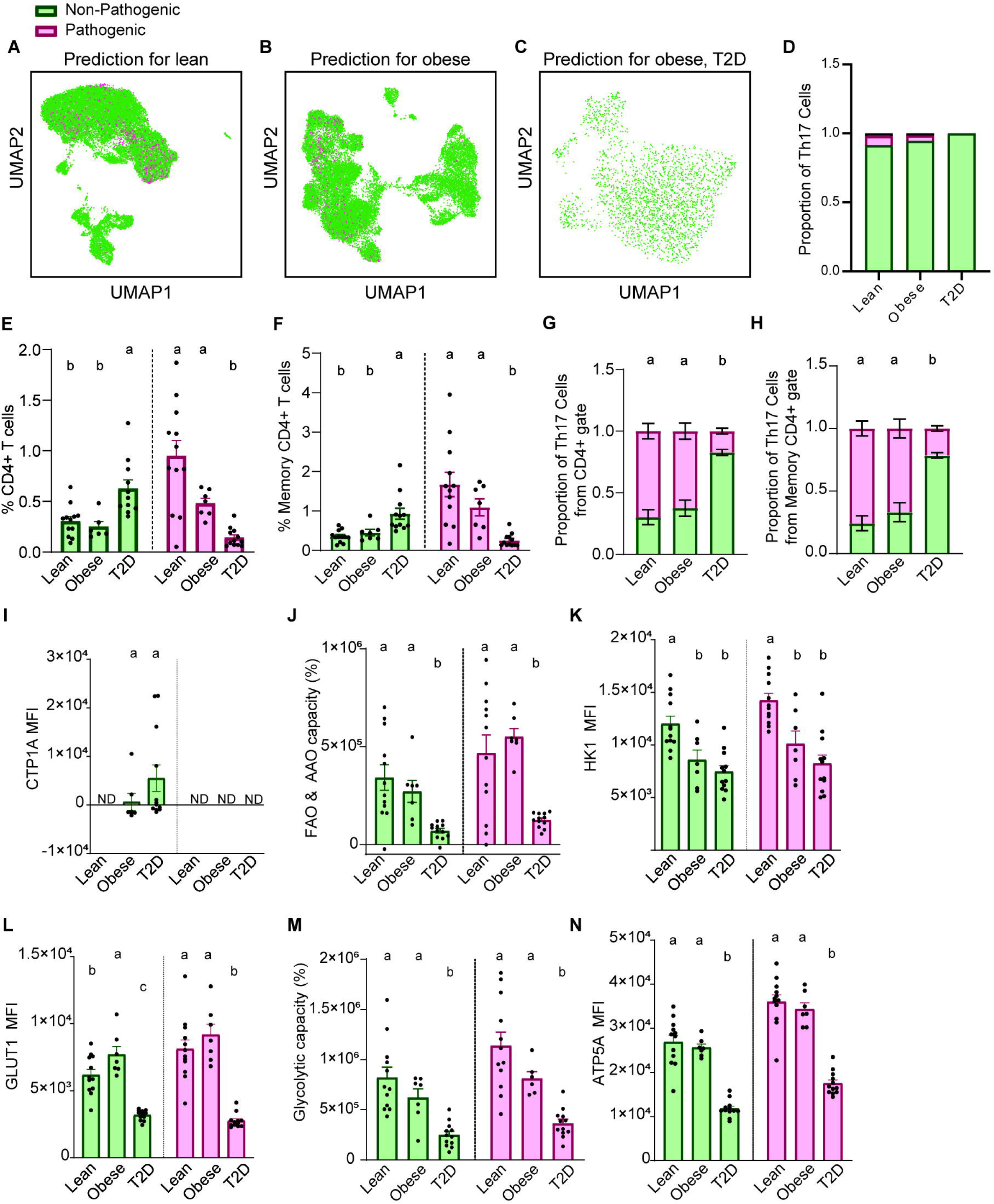
Th17np cells predominate in type 2 diabetes and exhibit impaired fatty acid oxidation despite elevated CPT1A. (**A to C**) UMAP plots showing transcriptomically classified Th17np (green) and Th17p (magenta) cells in published adipose tissue datasets from lean (A), obese (B), and obese type 2 diabetes (T2D) donors (C). (**D**) Proportions of classified Th17np and Th17p cells in the adipose tissue datasets. (**E and F**) Frequencies of Th17np and Th17p cells among total CD4+ T cells (E) and memory CD4+ T cells (F) from lean, obese, and obese T2D donors. (**G and H**) Proportions of Th17np and Th17p cells among total CD4+ T cells (G) and memory CD4+ T cells (H). (**I**) CPT1A median fluorescence intensity (MFI). (**J**) SCENITH-derived fatty acid oxidation and amino acid oxidation (FAO and AAO) capacity. (**K and L**) HK1 (K) and GLUT1 (L) MFI. (**M**) SCENITH-derived glycolytic capacity. (**N**) ATP5A MFI. In (E to N), green and magenta indicate Th17np and Th17p cells, respectively; bars show mean ± SEM, and points represent individual donors. Comparisons among lean, obese, and T2D groups were performed within each Th17 state (ANOVA). Different letters denote statistically significant groups (P < 0.05); groups sharing a letter are not significantly different. ND, not detected.

We observed a larger proportion of Th17np cells than Th17p cells in T2D donors in both the CD4+ and Memory CD4+ compartments when compared to lean and obese donors **(Fig 5E-H)**. Further analysis revealed that Th17np cells, specifically from T2D donors, exhibited high expression of CPT1A, but low FAO capacity when compared to lean and obese samples **(Fig 5I-J)**. Similar T2D-associated effects were observed in glycolytic capacity and glycolysis-associated enzyme expression. Th17np and Th17p cells from T2D donors had lower expression of HK1, GLUT1 and overall glycolytic capacity compared to lean and obese donors **(Fig 5K-M)**. Th17np and Th17p cells from T2D donors also differed in expression of ATP5A compared to lean and obese donors, consistent with our prior report of reduced ATP production in T cell stimulated PBMCs from T2D donors^43^ **(Fig 5N)**. PRDX2 expression was statistically lower only in Th17p T2D donors compared to lean and obese donors. Other metabolic enzymes (IDH2, G6PD, and ACAC) were measured but did not reveal T2D specific Th17np versus Th17p differences in metabolic phenotypes **(Supplemental Fig. 7A-D)**. Irrespective of T2D status, Th17p cells exhibited greater bioenergetic activity than Th17np cells, as supported by increased ATP5A expression and higher metabolic capacity **(Supplemental Fig. 7E-J).** Having identified impaired fatty-acid oxidation in T2D Th17np cells, we next asked whether broader changes in cellular lipid composition were detectable in T cells from the same metabolic environment. Untargeted lipidomics of bulk CD4⁺ T cells distinguished donors with T2D from metabolically healthy obese donors, with long-chain phosphatidylethanolamine, phosphatidylglycerol, and cardiolipin species contributing prominently to group separation **(Supplemental Fig. 8A-C)**. These data provide additional support of altered mitochondrial lipid remodeling in T2D T cells. Taken together, metabolic profiling of Th17 cells support the conclusion that Th17 cells from T2D are skewed toward Th17np cells, and that T2D-specific metabolic signatures involve similar metabolic programs predicted by COMPASS to be associated with Th17np cells.

### Acylcarnitine metabolism maintains human Th17np cells and creates a selective vulnerability in type 2 diabetes

Our transcriptomic analyses and metabolic phenotyping collectively implicated distinct metabolic pathways in the regulation of human Th17 state balance. To test whether these predicted pathways were functionally required for maintaining Th17np or Th17p cells, we performed a targeted CRISPR screen in primary human memory CD4⁺ T cells. Candidate genes were selected from Th17 state transcriptional signatures, COMPASS predictions, and metabolic profiling and included transcriptional regulators, glycosylation enzymes, and components of mitochondrial fatty-acid and acylcarnitine metabolism **(Fig. 6A, Supplemental Fig. 9A)**. CRISPR delivery and screen performance were evaluated by measuring viability, fluorescent dextran uptake, and cell recovery after activation **(Supplemental Fig. 9B-E)**. Knockdown of *IFNG* reduced IFN-γ expression, while Knockdown of *TCF7*, an established regulator of homeostatic Th17 identity, increased the relative representation of Th17p cells, consistent with its reported role in restraining proinflammatory RORγt activity^21^ **(Supplemental Fig. 9F-H)**. The screen identified CPT1A and the carnitine acylcarnitine translocase (CACT) as regulators of Th17 state balance **(Fig. 6A)**. CPT1A converts long-chain fatty acyl-CoAs into acylcarnitines, whereas CACT transports these acylcarnitines across the inner mitochondrial membrane (**Fig. 6D**). Kockdown of CPT1a resulted in expansion of the Th17p compartment. Conversely, knockdown of CACT resulted in expansion of Th17n cells (**Fig. 6B-C**). These data indicate that mitochondrial acylcarnitine transport may contribute to maintenance of Th17np states.

**Figure 6.**
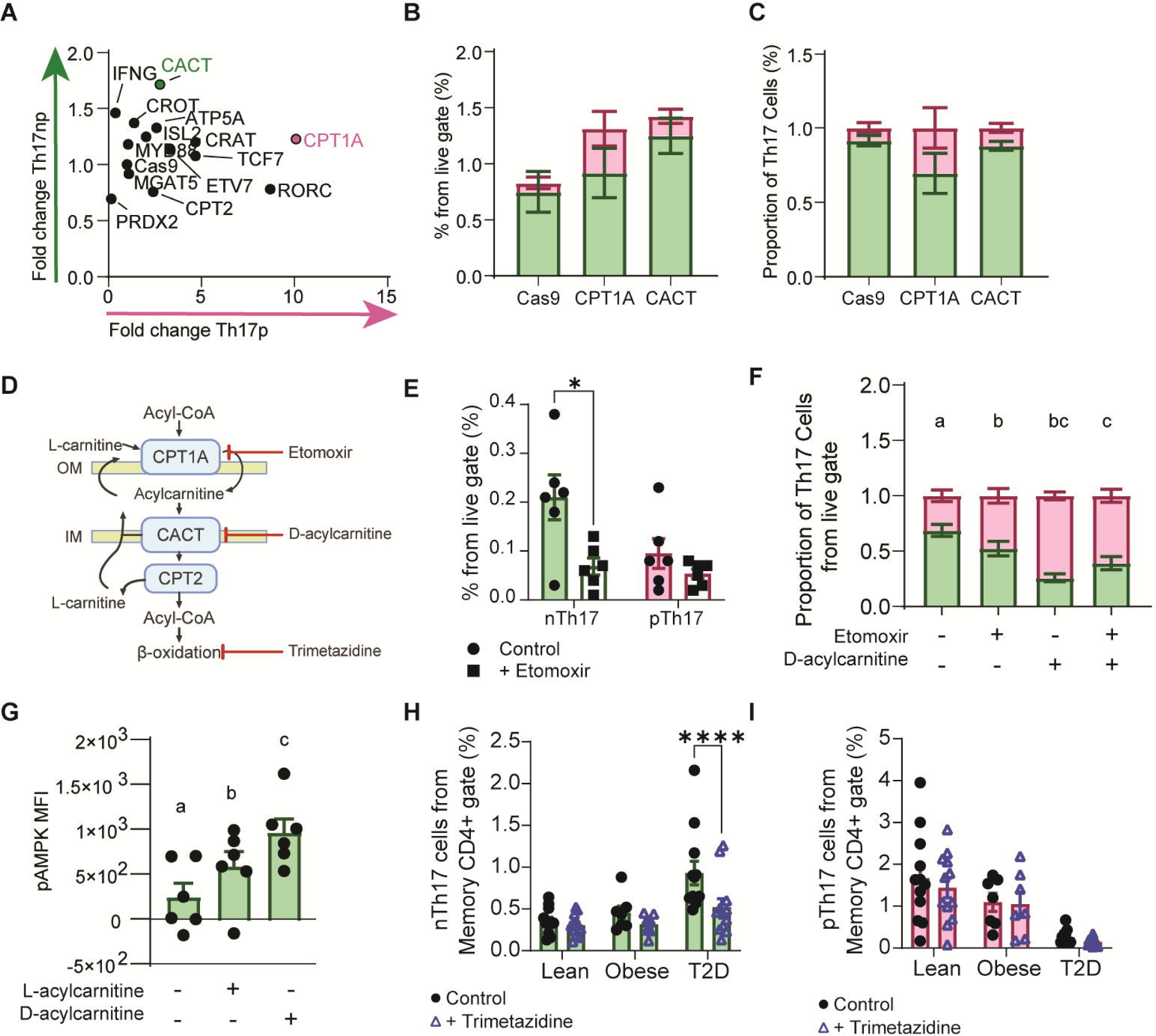
Acylcarnitine metabolism maintains human Th17np cells and creates a selective vulnerability in type 2 diabetes. (**A**) Targeted CRISPR screen showing fold changes in Th17np and Th17p frequencies after knockdown of the indicated genes relative to the Cas9 only control. (**B**) Frequencies and (**C**) proportions of Th17np and Th17p cells among live cells after Cas9 only control, CPT1A knockdown, or CACT knockdown. (**D**) Schematic of the mitochondrial carnitine shuttle and sites of pharmacologic inhibition. CPT1A (carnitine palmitoyltransferase 1A); CACT (carnitine-acylcarnitine translocase); OM, (outer mitochondrial membrane); IM, (inner mitochondrial membrane). (**E**) Frequencies of Th17np and Th17p cells among live cells from healthy lean donors treated with vehicle or etomoxir. (**F**) Proportions of Th17np and Th17p cells after treatment with vehicle, etomoxir, D-acylcarnitine, or etomoxir plus D-acylcarnitine. (**G**) Phosphorylated AMPK (pAMPK) MFI after treatment with vehicle, L-acylcarnitine, or D-acylcarnitine. (H and I) Frequencies of Th17np (H) and Th17p (I) cells among memory CD4+ T cells from lean, obese, and obese T2D donors treated with vehicle or trimetazidine. Bars show mean ± SEM; points represent independent donors. In (F) and (G), different letters denote statistically significant groups (P < 0.05); groups sharing a letter are not significantly different. *P < 0.05 and ****P < 0.0001.

We next tested the impact of pharmacological inhibition of CPT1A and CACT on Th17 state balance. In memory CD4⁺ T cells from healthy lean donors, inhibition of CPT1A with etomoxir selectively reduced Th17np cells (**Fig. 6E**). The unnatural D isomer of long-chain acylcarnitines strongly inhibits CACT (Fig. 6D)^46^, but can also induce toxicity from accumulation of acylcarnitines. We tested the short-term impact of etomoxir or exogenous D-acylcarnitine on Th17 state balance and found that D-acylcarnitine, similar to etomoxir reduces Th17np cells, though selective cell death cannot be excluded (**Fig. 6F, Supplemental Fig. 9I**). These data suggest that generation of acylcarnitine and acylcarnitine abundance influence Th17 state maintenance. Because AMPK signaling has been associated with suppression of pathogenic Th17 programs, we tested whether exogenous long chain acylcarnitine induced AMPK activation. We found that both L and D isomers of acylcarnitine increased AMPK phosphorylation in Th17np cells. These data link increased acylcarnitine availability to known mechanisms of Th17p suppression (**Fig. 6G**).

Because T2D Th17np cells exhibited elevated CPT1A expression concurrent with impaired fatty-acid oxidation capacity, we next asked whether this discordant metabolic state created a disease-selective vulnerability. Using the fatty-acid oxidation inhibitor trimetazidine to reduce overall lipid flux and shift cellular metabolism toward glycolysis selectively reduced Th17np cells from donors with T2D, whereas healthy donor Th17np cells and Th17p cells from both groups were unaffected (**Fig. 6H-I**). Together, these results identify mitochondrial acylcarnitine metabolism as a regulator of human Th17np cell maintenance and indicate that human Th17np cell maintenance depends on acylcarnitine flux rather than simply increased fatty acid uptake or oxidation. Thus, the impaired oxidative state of T2D Th17np cells creates a selective metabolic dependency that distinguishes this population and provides a potential strategy for its targeted modulation.

## Discussion

The inability to prospectively identify Th17 cell states in primary human cells has limited efforts to determine whether the transcriptional and metabolic programs defined in murine models apply to human disease. Here, we established complementary single cell transcriptomic and flow cytometry frameworks for annotating human Th17 states directly from primary samples. Application of these tools across tissues and diseases identified transcriptional features conserved with murine Th17 programs, revealed distinct bioenergetic characteristics of human Th17 subsets, and uncovered an unexpected predominance and metabolic vulnerability of Th17np cells in T2D. Collectively, these findings provide a framework for studying human Th17 heterogeneity while demonstrating that the historical labels “pathogenic” and “nonpathogenic” do not necessarily predict which state predominates in a given human disease.

Our nanovial scELISA enabled enrichment of viable, cytokine-secreting Th17 cells for transcriptional analysis without relying on *in vitro* polarization of naïve T cells. While resource-intensive (∼8 million input cells yielding ∼2,000 enriched Th17 cells), it provides a discovery platform for characterizing rare primary populations that are otherwise diluted within heterogeneous PBMC cultures. The resulting transcriptional signatures enabled classification of Th17 states in larger single cell datasets and informed a scalable flow cytometry gating strategy. Our classification model allows transcriptomic discovery using existing and newly generated single cell datasets, while the flow cytometry gating strategy enables population level assessment across clinical cohorts, thereby democratizing study of Th17 subsets in humans.

The human Th17 states identified by this framework retained core features of murine Th17np and Th17p programs across multiple sample sources. Human Th17p cells displayed immune-activation pathways and expression of cytokines associated with murine pathogenicity^18–20^, whereas ETV7 and ISL2 emerged as reproducible features of the human Th17p state. Their functions in Th17 biology remain unknown, but their conservation across tissue and disease contexts nominates them as candidate regulators or markers of this state. Similarly, increased complex N-glycan branching in Th17p cells provided functional support for the metabolic predictions generated by COMPASS. These data are consistent with N-glycan branching supporting the fate decision Th17p cells and Tregs^47^. Whether glycan branching is required to establish Th17p identity or instead reflects their activation and metabolic state will require direct perturbation of the responsible enzymes.

Th17np cells in T2D exhibit high CPT1A expression alongside paradoxically reduced FAO capacity, consistent with disrupted mitochondrial lipid metabolism previously reported in T2D PBMCs^43^. Impaired downstream oxidation may drive cytosolic acylcarnitine accumulation, where acylcarnitines function as signaling molecules rather than metabolic intermediates. Increased delivery of fatty acids into the carnitine shuttle without commensurate downstream oxidation could disrupt acylcarnitine homeostasis, activate energy-stress signaling, and create dependence on residual mitochondrial lipid flux. Consistent with this model, T2D Th17np cells were selectively susceptible to trimetazidine, whereas Th17p cells were comparatively resistant. Impaired fatty-acid oxidation may therefore be more than a descriptive feature of T2D Th17np cells. Direct measurement of acylcarnitine species in sorted Th17 states and isotope-tracing experiments will be necessary to establish the direction and magnitude of pathway flux. However, the reliance on intracellular cytokine staining to identify Th17 cell states currently precludes live cell downstream assays on Th17np vs Th17p cells. Future identification of surface markers from our transcriptomic datasets may permit prospective isolation and functional testing.

Finally, these findings challenge whether the terms “pathogenic” and “nonpathogenic” are the most accurate descriptors of Th17 subtypes. “Pathogenic” and “nonpathogenic” were defined largely through cytokine polarization and adoptive transfer experiments in murine autoimmune models, in which pathogenicity could be established by the capacity of a population to cause disease^33,48–50^. In human studies, however, these terms are commonly assigned from transcriptional resemblance, cytokine production, or the inflammatory setting in which cells are found, rather than from a direct test of pathogenic function^51–54^. Our finding that Th17np classified cells predominate in T2D illustrates this disconnect. This finding is further supported by a lack of identification of IFN-γ producing Th17 cells in T2D^55^. In our data, a population described as “nonpathogenic” can be expanded within a chronic inflammatory disease. Therefore, a population bearing the canonical “pathogenic” signature does not inherently dominate every inflammatory condition. Conversely, our data demonstrate that the designation “nonpathogenic” should not be interpreted as uniformly protective in human disease. We therefore suggest that the field should revisit the use of pathogenicity as a fixed identity for human Th17 cells^56^.

In summary, this study establishes an integrated framework for identifying and functionally interrogating human Th17 states. Application of this framework reveals that the Th17 state historically designated “nonpathogenic” predominates in T2D and acquires a selective dependence on mitochondrial lipid metabolism. These findings support revisiting nomenclature in which human Th17 states are defined pathogenic labels inherited from murine models. More broadly, this work establishes that the metabolic liabilities of human Th17 cells are disease-context dependent, creating opportunities to modulate maladaptive populations without treating the entire Th17 compartment as a uniform therapeutic target.

## Materials and Methods

### Biological samples

#### Human blood, PBMC, and tonsil samples

PBMC samples from donors with multiple sclerosis (n = 8) were obtained in collaboration with Michael Sy at UCI under IRB protocol 1496 (**Supplemental Table 1**). Samples from donors with asthma (n = 3) were obtained in collaboration with Dawn Newcomb at Vanderbilt University Medical Center (VUMC) under IRB protocol 202162. All participants provided written informed consent, and all procedures were conducted in accordance with the Declaration of Helsinki and applicable institutional guidelines.

Whole blood from normoglycemic lean (body mass index [BMI] ≤28 kg/m²; n = 12) and normoglycemic obese (BMI >28 kg/m²; n = 7) donors was obtained through the University of California, Irvine Institute for Clinical and Translational Science (UCI ICTS). Cryopreserved PBMCs from additional lean donors were obtained from Sanguine Biosciences, and cryopreserved PBMCs from obese donors with type 2 diabetes (T2D; n = 12) were obtained from Charles River Laboratories. Donor group definitions and available clinical characteristics are provided in **Supplemental Table 2**.

Independent single-cell RNA-sequencing datasets from matched peripheral blood and tonsil samples were analyzed as previously described^7^. Published adipose-tissue single-cell datasets from lean, obese, and T2D donors were analyzed as described in references 9 and 10.

#### Experimental autoimmune encephalomyelitis model

EAE was induced in 17-week old male C57BL/6 mice via subcutaneous injection of 150uL of MOG_35–55_ peptide emulsified in complete Freund’s adjuvant (CFA) supplemented with heat-killed Mycobacterium tuberculosis (Hooke Laboratories EK-0111) on the hindquarters of the animal. Two injections of 200uL (1ug/mL) of pertussis toxin were given to each animal at 2 and 48 hours via intraperitoneal injection for a total of 400uL. Inguinal lymph nodes (iLN) were collected and mechanically dissociated with a 50um filter and pestle and washed with FACs buffer, then resuspended at 1.25×10^6^/mL with R10 media and 200uL seeded in a flat-bottom 96-well plate for activation and flow cytometry.

### CD4+ T-cell enrichment and culture

#### Memory and naïve CD4+ T-cell enrichment

Memory (130-091-893) and naïve (130-094-131) CD4+ T cells were enriched from PBMCs using Miltenyi Biotec’s negative isolation kits. PBMCs were quickly thawed in a 37C water bath and washed with 10mL of RPMI 1640 media (Genesee Scientific, CAT#25-506) supplemented with 10% FBS (Omega Scientific, FB-1), 20uM HEPES (ThermoFisher Scientific, 15630080), 100ug/mL Penicillin/Streptomycin (Genesee, 25-512).

#### In vitro differentiation of Th17np and Th17p cells

Human naive CD4+ T cells were cultured in 37°C 5% CO_2_ incubator for 5 days in anti-CD3 (1:1,000) (ThermoFisher, CAT# 344814) coated 96 well U-bottom plates in the presence of IL-6 (Miltenyi Biotec, CAT#130095365), TGFꞵ (Miltenyi Biotec, CAT#130095067) and anti-CD28 (ThermoFisher, CAT#14-0289-82) for non-pathogenic Th17 cells, or IL-1ꞵ (Miltenyi Biotec, CAT#130093895), IL-6 (Miltenyi Biotec, CAT#130095365), IL-23 (1:500) (Miltenyi Biotec, CAT#130095757) and anti-CD28 (ThermoFisher, CAT#14-0289-82) pathogenic Th17 cells.

#### CD4+ T-cell activation

96 well U-bottom plates were coated with anti-CD3 (1:1,000) (ThermoFisher, CAT#344814). 200uL of cells were then seeded at 1.25×10^6^/mL with R10 media and anti-CD28 (1:500) (ThermoFisher, CAT#14-0289-82) for 40 hours in 5% CO_2_ incubator. Last 5 hours cells were supplemented with brefeldin-A (1:1,000) (Biolegend, CAT#420601) and then harvested and stained for flow cytometry.

### Nanovial single-cell cytokine-capture assay

Biotinylated nanovials (Partillion Biosciences, NV135BT-01) were functionalized sequentially with streptavidin (Thermo Fisher Scientific, 434302) and biotinylated antibodies against human CD45 (BioLegend, 304004), IL-17A (PeproTech, 500-P07Gbt), IL-17F (PeproTech, 500-P90bt), and IL-21 (PeproTech, 500-P191bt). Activated memory CD4+ T cells were combined with functionalized nanovials at a 4:1 nanovial-to-cell ratio to achieve no more than one cell per nanovial. Unbound cells were removed by filtration through a 25-µm strainer, which retained the 35-µm nanovials while allowing free T cells to pass. Cell-loaded nanovials were incubated for 4 h at 37°C and 5% CO₂ to capture locally secreted cytokines.

Secreted cytokines were detected using Core Quantum-conjugated fluorescent magnetic secondary antibodies: MagDot 575 anti-IL-17A (PeproTech, 500-P07), MagDot 610 anti-IL-17F (PeproTech, 500-P90), and MagDot 640 anti-IL-21 (PeproTech, 500-P191). Cytokine-positive nanovials were enriched magnetically, and the recovered viable Th17 cells were used for flow cytometry and single-cell RNA sequencing. Approximately 8 × 10⁶ input cells yielded approximately 2,000 enriched Th17 cells; cytokine-positive cells constituted 90.93% of viable recovered cells.

### Single-cell RNA sequencing

Single-cell suspensions were loaded on a Chromium X controller (10X Genomics) with a loading target of 30,000 - 50,000 (depending on the sort yield) with a goal of sequencing ∼500,000 cells per donor. Gene expression libraries were generated using the Chromium Next Gem Single Cell 5’ Reagent Kit v2 (Dual Index) per the manufacturer’s instructions. Quality and quantity of libraries were measured on tapestation, qubit, and bioanalyzer and sequenced on Illumina NovaSeq 6000 with a sequencing target of 30000 reads per cell for gene expression libraries.

### Single-cell transcriptomic analysis

#### Quality control, integration, and cell annotation

Raw 10x Genomics data from in vitro-skewed Th17np and Th17p cells, multiplexed samples, and nanovial-enriched ex vivo samples were processed using a standardized quality-control pipeline. Ambient RNA was removed with CellBender, doublets were identified with DoubletDetection, and cells were retained if they expressed at least 200 genes, contained no more than 25% mitochondrial transcripts, and fell within ±5 median absolute deviations of sample-level quality-control distributions. CD4+CD161+ cells were annotated with CellTypist immune-reference models.

Before integration, sample-associated batch structure was quantified (R²sample = 0.20; entropy = 0.14). scVI was trained for 200 epochs on 8,000 highly variable genes supplemented with Th17 marker genes using a 20-dimensional latent space. Integration reduced sample-associated structure (R²sample = 0.07; entropy = 0.31).

#### Neural network

Raw 10×Genomics single-cell RNA sequencing data from *in vitro* skewed pathogenic (P) and non-pathogenic (NP) Th17 cells, multiplexed samples, and nanovial-enriched, *ex vivo* donor samples were processed through a standardized quality control pipeline comprising ambient RNA removal (CellBender), doublet detection (DoubletDetection), and low-quality cell exclusion based on minimum gene detection (≥200 genes), mitochondrial RNA content (≤25%), and median absolute deviation-based outlier removal (±5 MAD). CD4+CD161+ cells underwent cell type annotation using CellTypist with immune reference models, and pre-integration batch assessment revealed substantial technical batch effects (R²_sample = 0.20, entropy = 0.14), which were corrected using scVI trained on 8,000 highly variable genes supplemented with Th17 marker genes over 200 epochs in a 20-dimensional latent space, successfully harmonizing data across donors and experimental conditions (R²_sample = 0.07, entropy = 0.31). Pathogenicity classification was subsequently performed using SCANVI, a semi-supervised deep generative model that jointly leverages labeled *in vitro* P and NP cells and unlabeled *ex vivo* cells, evaluated via 5-fold stratified out-of-fold cross-validation with a decision rule of probability threshold 0.5 and confidence margin Δ = 0.30, achieving an accuracy of 0.94, precision of 0.92, recall of 0.94, F1 score of 0.93, and approximately 2% ambiguous cells; permutation testing (AUC ≈ 0.5) confirmed genuine biological signal. The final model was applied to independent peripheral blood and tonsil datasets from collaborating institutions using scArches-based transfer learning, yielding consistent and biologically plausible classifications across tissue types and donor cohorts, with ambiguous cell rates of 4.2% and 2.0% respectively.

Differential expression (DE) analysis was performed using the built-in differential_expression method of the scVI/SCANVI framework (mode=’change’, weights=’importance’), which accounts for learned latent structure and uncertainty in cell state assignments. DE genes were identified using stringent criteria of Bayes factor ≥ 3 and |log₂ fold-change| ≥ 1, and results were visualized using volcano plots. Cross-dataset overlap of DE genes across in vitro, ex vivo, peripheral blood, and tonsil datasets were assessed using UpSet plots to identify genes consistently differentially expressed independent of tissue source or experimental context. Gene set enrichment analysis (GSEA) and over-representation analysis (ORA) were subsequently performed using MSigDB v2025.1 to identify biological pathways enriched among genes upregulated in pathogenic (Up_in_P) or non-pathogenic (Up_in_NP) cell states, with results compiled across gene set collections and visualized as per-GMT barplots. Pathways and DE genes reaching significance across all four datasets were identified and visualized using cross-dataset heatmaps.

#### COMPASS metabolic modeling

Single-cell metabolic states were inferred using COMPASS (Wagner et al., 2021), a constraint-based framework that integrates normalized single-cell transcriptomic data with flux balance analysis (FBA) to generate cell-by-reaction penalty scores. COMPASS accepts annotated data sets (h5ad) as inputs and produces a tab-separated values (TSV) output file (reactions.tsv) output matrix. During postprocessing, these raw penalties are transformed into reaction consistency scores, where higher values indicate greater predicted reaction activity. In our study, cells classified as Th17p or Th17np were analyzed separately within each source dataset, and COMPASS was run independently on SCANVI classified Th17p and Th17np cells from in vitro, ex vivo, blood, and tonsil datasets using the “homo_sapiens” setting for species.

For downstream analysis, the annotated datasets were imported into R (version 4.5.2) to generate cell-metadata tables and linear gene-expression matrices, which were reordered to match the cell order in the corresponding COMPASS output files. Using the compassR package (version 1.0.0), reaction consistency matrices and associated reaction metadata were loaded for each Th17p and Th17np run, summarized at the subsystem level, and used to compute per-cell pathway activity scores. For direct Th17p versus Th17np comparisons within each dataset, shared reactions were aligned between paired runs and tested using unpaired Wilcoxon rank-sum tests with Cohen’s d effect sizes, following the COMPASS postprocessing framework. Reactions were then annotated with subsystem, Enzyme Commission (EC) number, and confidence metadata and aggregated to pathway-level summaries. Figures generated in R for these analyses included ranked subsystem bar plots and volcano plots.

### Flow Cytometry

Cells were first stained with live/dead fixable dye diluted in PBS, pH 7.4 for 20 minutes on ice away from light. CD4+ T cells were washed with FACs buffer and centrifuged as 500g for 5 minutes. Supernatant was removed and 25uL of Human TruStain FcX was added for 10 minutes on ice away from light to block human FC receptor, then washed, centrifuged and supernatant removed. Then 25uL of surface antibody master mix diluted in BD Biosciences Brilliant Stain Buffer (CAT# 566349) was added to CD4+ T cells for 1 hour on ice away from light. CD4+ T cells were then washed with FACs buffer, centrifuged and supernatant removed. CD4+ T cells were then fixed with ThermoFisher’s Foxp3 Fixation Buffer (00-5523-00) for 30 minutes at room temperature away from light. CD4+ T cells were washed with ThermoFisher’s Foxp3 Intracellular Staining Permeabilization Wash Buffer (00-5523-00) twice and supernatant was removed for the addition of 25uL of intracellular antibody master mix diluted in ThermoFisher’s Foxp3 Intracellular Staining Permeabilization Wash Buffer overnight in 4℃ away from light. CD4+ T cells were washed with ThermoFisher’s Foxp3 Intracellular Staining Permeabilization Wash Buffer and resuspended in 200uL of 1% paraformaldehyde diluted in PBS, pH 7.4 for the performance of spectral flow cytometry on Cytek’s 3-laser Northern Lights.

#### Controls

Antibodies were titrated and a dilution was chosen based on optimal resolution discriminating between high and low/negative fluorescence intensity. Single stain controls were performed on CD4+ T cells or Thermo Fisher’s Ultra-comp PLUS beads (CAT#01-3333-42) depending on fluorescence intensity and marker expression. Fluorescence minus one (FMO) controls were produced to set accurate gates on particularly dim and rare markers. Unmixing was performed on SpectroFlo software using single-stained controls and unstained cell sample to account for autofluorescence.

#### L-PHA staining

Cells were treated with 10uM kifunensine at time of activation and then proceeded with flow cytometry protocol outline above.

#### SCENITH and Metflow

Flow cytometry panel design was inspired by Arguello et al. (2020) and Ahl et al. (2020), where SCENITH measured protein translation as a proxy to measure metabolic flux. Memory CD4+ T cells were treated with DMSO (1:1000) (MiliporeSigma, CAT#D2650-5X5ML), 100mM 2DG (Sigma-Aldrich, CAT#D8375-5G), 1.5uM Oligomycin (Fisher Scientific, CAT#49545510MG), 10uM CB-839 (MedChemExpress, CAT#HY-12248), 5uM Trimetazidine (MedChemExpress, CAT#HY-B0968A), or a combination of 2DG and Oligomycin for 30 minutes along with brefeldin-A in 37°C 5% CO_2_. After 30 minutes, cells were treated with 10uM puromycin for 40 minutes in 37°C 5% CO_2_ and then washed with PBS for flow cytometry staining protocol.

Met-flow consisted of nine metabolic proteins chosen based on their critical role in specific metabolic pathways. The following metabolic antibodies were purchased from Abcam: SLC20A1, HK1, CPT1A, IDH2, G6PD, ASS1, and PRDX2, and custom conjugated in-house to their respective fluorophores using various conjugation kits. Negative MFIs after unmixing were denoted ND.

#### Untargeted Lipidomics

CD4+ T cells from normoglycemic obese (n = 5) and obese T2D (n = 5) donors from the previously described cohort^43^ were activated for 40 h and enriched using anti-CD3 Dynabeads before submission to the Research Mass Spectrometry and Proteomics Core (RMSPC) for untargeted lipidomic analysis. Lipid extraction, liquid chromatography mass spectrometry acquisition, feature identification, normalization, and quality control were performed using established core procedures. Lipid species associated with mitochondrial membranes were selected for downstream analysis. Partial least-squares discriminant analysis (PLS-DA) and variable-importance-in-projection analyses were performed in an R Shiny workflow in the CytokineProfile integrated web tool^57^.

#### CRISPR–Cas9 Knockdown

Memory CD4+ T cells from three lean donors were rapidly thawed and washed in supplemented RPMI 1640 as described above. Ribonucleoprotein complexes containing Alt-R HiFi Cas9 and target-specific single-guide RNAs (sgRNAs; **Supplemental Table 4**) were delivered using the Portal Biotechnologies Gateway mechanoporation platform with an 8-psi setting and a 5-µm-pore chip. For each condition, approximately 6 × 10⁵ cells in 50 µL were combined with Cas9, 3 pooled sgRNA, basal medium, and 10× PBS according to the manufacturer’s protocol. Cas9-only and no-mechanoporation controls were included. Cells were subsequently activated for 40 h, treated with brefeldin A during the final 5 h, and analyzed by flow cytometry.

### Statistical Analysis

Statistical analyses were performed in R 4.2.3 and GraphPad Prism 9.5.1. Continuous data were assessed for normality using Shapiro–Wilk tests and Q–Q plots. Two-group comparisons were analyzed using two-tailed Student’s or Welch’s t tests when normality assumptions were met and Mann–Whitney U tests otherwise. A paired t test was used for IFN-γ knockdown relative to the matched Cas9 control. Proportions were compared using z tests. Experiments with more than two groups were analyzed by one-way analysis of variance followed by Dunnett’s multiple-comparisons test for normally distributed data or by Kruskal–Wallis tests for nonparametric data. Two-way analysis of variance was used for the L-PHA experiment. Comparisons among lean, obese, and T2D cohorts in the primary metabolic analysis were performed using Kruskal–Wallis tests; one-way analysis of variance was used for the supplemental cohort comparisons within each Th17 state. COMPASS and transcriptomic analyses used the procedures described above. P < 0.05 or adjusted P < 0.05, as applicable, was considered significant. Bars show mean ± s.e.m. unless otherwise indicated, and each point represents an independent donor or biological replicate.

No statistical method was used to predetermine sample size. Samples were not randomized, and investigators were not blinded to experimental groups.

## Data and Code Availability

Analysis code is available at https://github.com/surPoudel/NP_P_Th17_prediction_application. Newly generated single-cell RNA-sequencing data will be deposited in a public repository before publication. Published datasets were obtained from the sources cited in references (^44,45^). Additional data supporting the findings are available from the corresponding author upon reasonable request.

## Acknowledgements

We thank Dr. Mkhikian (University of California, Irvine) for providing the L-PHA reagent and guidance on lectin staining and Dr. Matthew Griffin (University of California, Irvine) for assistance with interpretation of the COMPASS analyses.

## Author contributions

Conceptualization: N.S.U., S.P., and D.N. Methodology: N.S.U., S.P., S.Sa., S.Su., J.K., U.-V.L., A.R.J., N.H., T.S., M.M., E.H.P., S.B., and C.W. Investigation: N.S.U., J.K., U.-V.L., A.R.J., N.H., T.S., M.M., E.H.P., S.B., and C.W. Software: S.P. Formal analysis: N.S.U., S.P., S.Sa., and D.N. Data curation: S.P., S.Su., M.Mej., S.B., and C.W. Visualization: N.S.U. Resources: S.Su., M.S., D.Nc., S.B., C.W., C.Wa., L.E.W., B.S.N., X.D.Z., D.G., and D.N. Funding acquisition: C.Wa., L.E.W., B.S.N., X.D.Z., D.G., and D.N. Supervision: C.Wa., L.E.W., B.S.N., X.D.Z., D.G., and D.N. Project administration: D.N. Writing—original draft: N.S.U., S.P., and D.N. Writing—review and editing: All authors. All authors contributed to interpretation and discussion of the findings and approved the final manuscript.

