## Supplemental Fig. for "Human Th17-state classification reveals a selective vulnerability in acylcarnitine metabolism in type 2 diabetes"

**Supplemental Figures 1-9**

**Supplemental Tables 1-2**

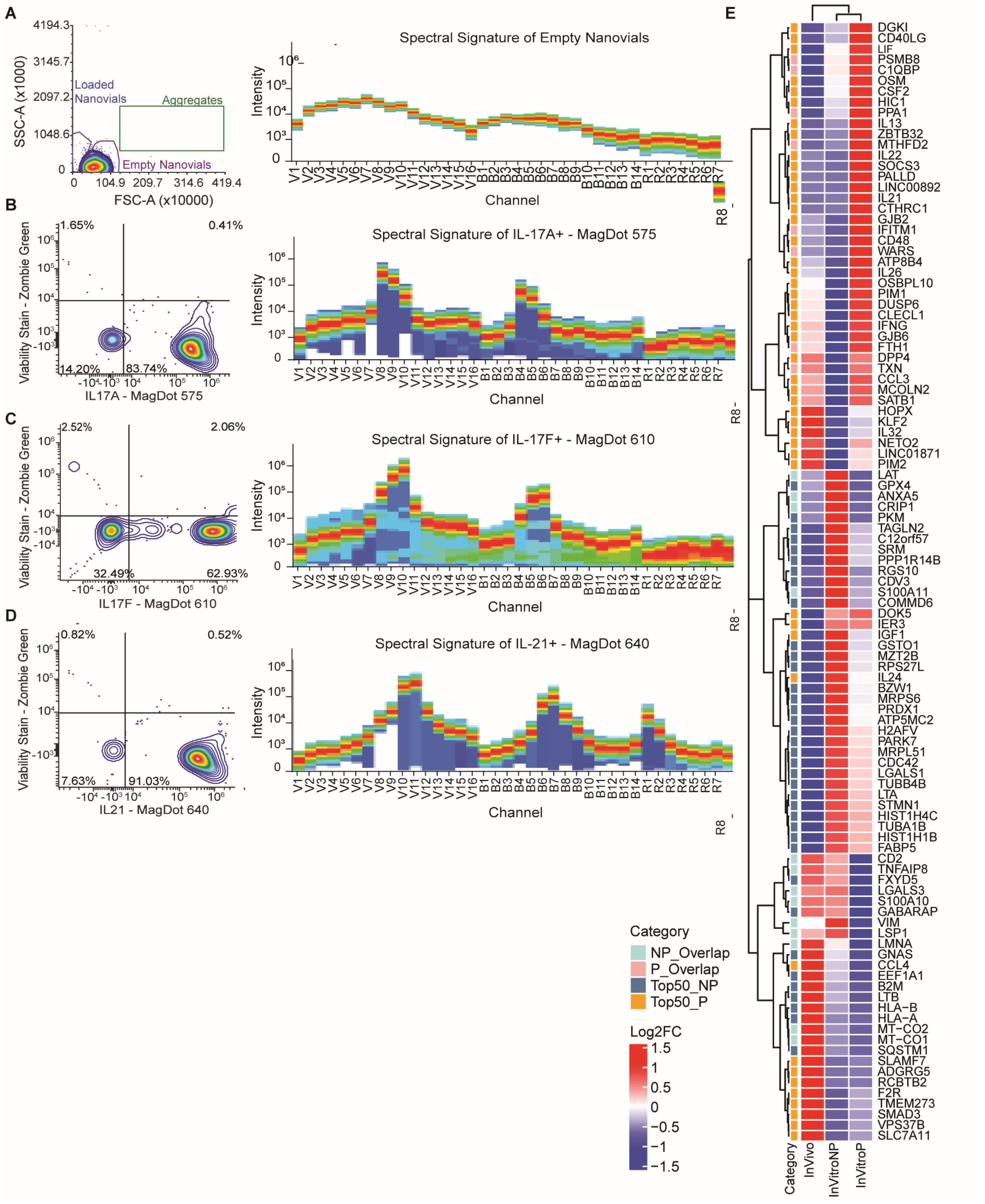


**Supplemental Figure 1. Validation of nanovial cytokine detection and transcriptional programs in enriched human Th17 cells.** (**A**) Representative flow cytometry discrimination of empty, cell-loaded, and aggregated nanovials and spectral signature of empty nanovials. (**B to D**) Representative viability-versus-cytokine plots and spectral signatures for IL-17A detected with MagDot 575 (B), IL-17F detected with MagDot 610 (C), and IL-21 detected with MagDot 640 (D). (**E**) Hierarchical clustering of differentially expressed genes across nanovial-enriched *ex vivo* and *in vitro*-skewed Th17np and Th17p cells. The annotation bar denotes genes shared between conditions or among the top 50 genes associated with either state.

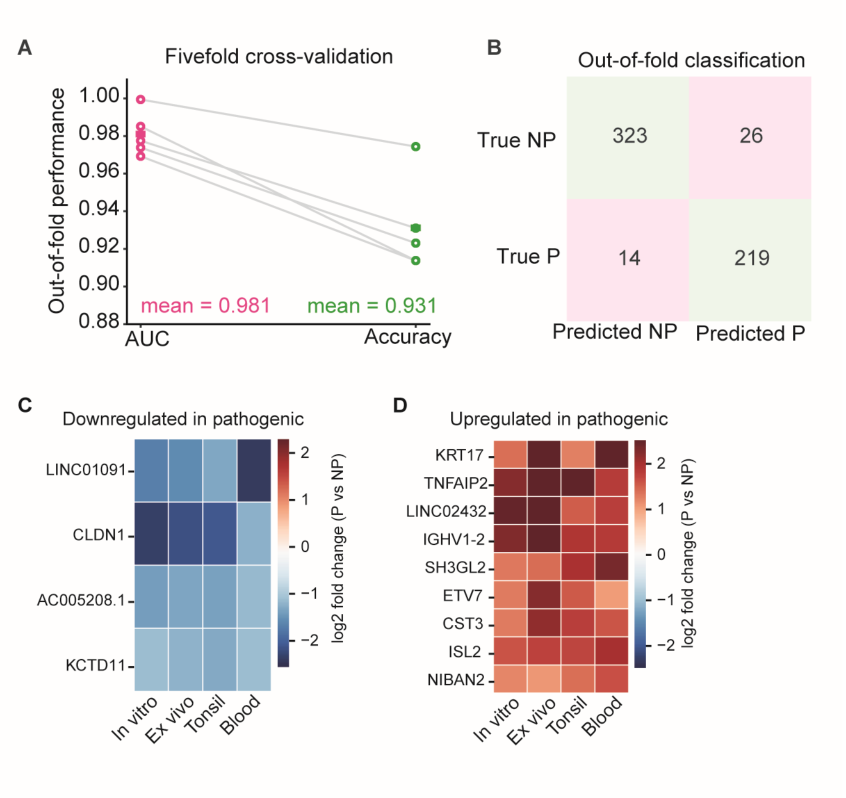


**Supplemental Figure 2. Cross-validation and molecular characterization of the Th17 state classifier.** (**A**) Fivefold cross-validation of the scANVI classifier showing out-of-fold area under the receiver operating characteristic curve (AUC) and accuracy for each fold. Mean values are indicated. (**B**) Out-of-fold confusion matrix showing true and predicted Th17np and Th17p classifications. (**C and D**) Heatmaps showing log2 fold changes in genes consistently down-regulated (C) or up-regulated (D) in Th17p relative to Th17np cells across *in vitro*, *ex vivo*, tonsil, and blood datasets.

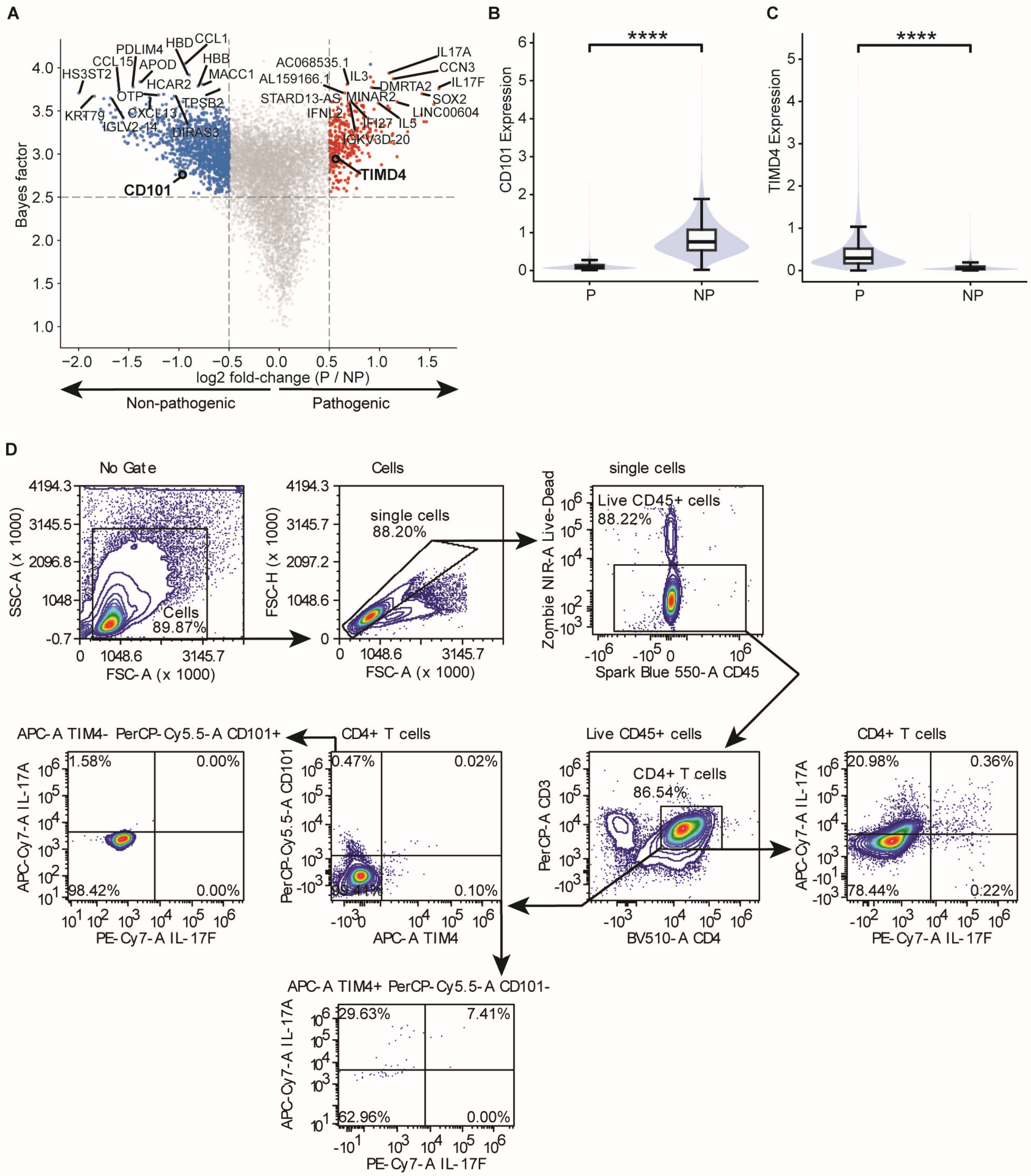


**Supplemental Figure 3. Candidate surface markers do not reliably identify primary human Th17 states.** (**A**) Differential gene-expression analysis of *in vitro*-skewed Th17p relative to Th17np cells. The x axis shows log2 fold change (Th17p/Th17np), and the y axis shows Bayes factor. Selected genes, including the surface marker candidates CD101 and TIMD4, are labeled. Dashed lines indicate the differential-expression thresholds. (**B and C**) Violin and box plots of CD101 (B) and TIMD4 (C) expression in transcriptomically classified Th17p and Th17np cells. (**D**) Representative flow cytometry strategy evaluating CD101 and TIM4 surface expression and IL-17A and IL-17F production in primary CD4+ T cells from a healthy lean donor. ****P < 0.0001.

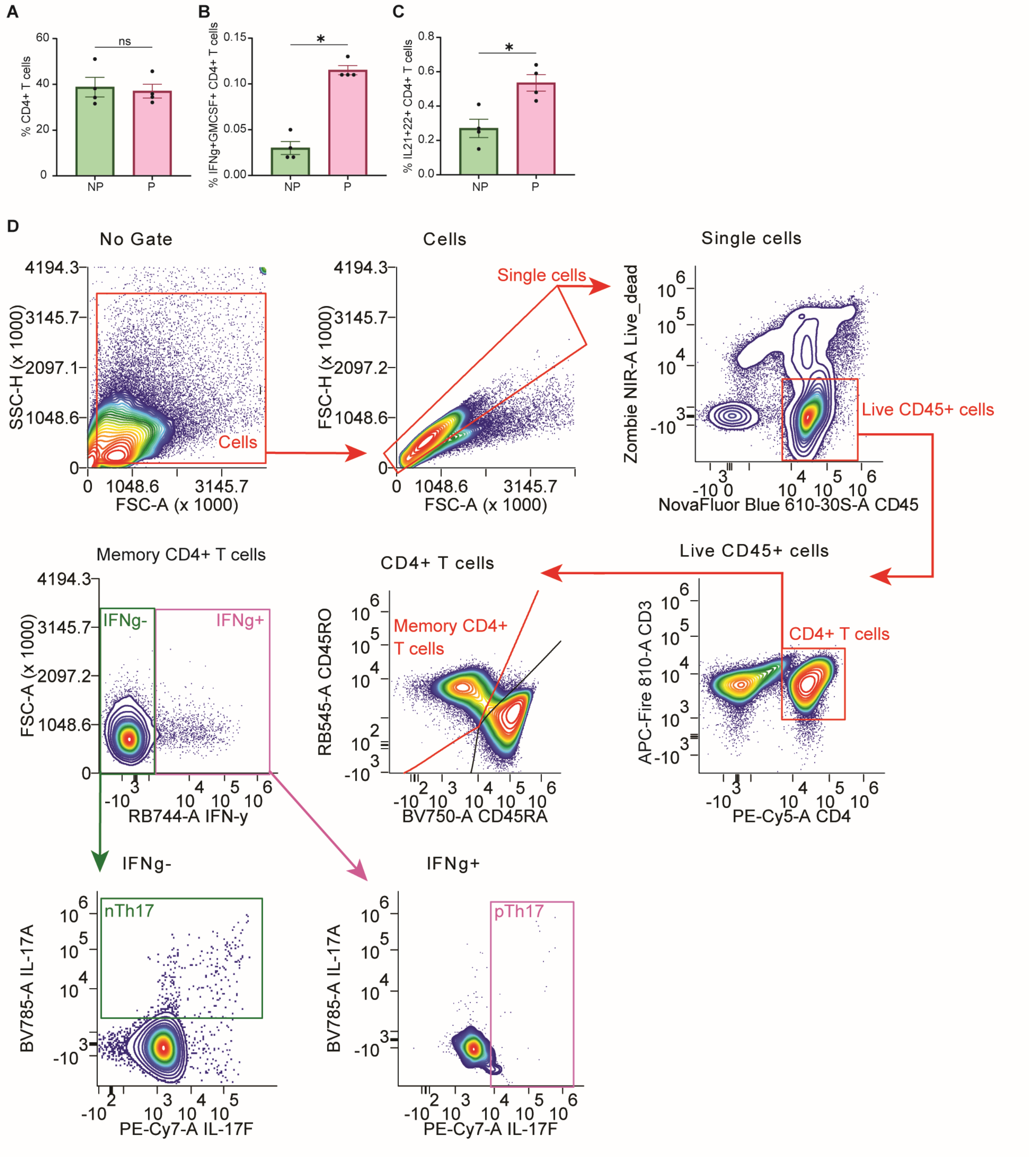


**Supplemental Figure 4. Validation of Th17p-associated cytokines and the Th17 state gating strategy.** (**A to C**) Frequencies of total CD4+ T cells (A), IFN-γ+GM-CSF+ CD4+ T cells (B), and IL-21+IL-22+ CD4+ T cells (C) under Th17np and Th17p skewing conditions (n = 4 donors). (**D**) Representative gating, singlets, live CD45+ cells, CD4+ T cells, memory CD4+ T cells, IFN-γ− and IFN-γ+ populations, and Th17np (IFN-γ−IL-17A+) and Th17p (IFN-γ+IL-17F+) cells. Bars show mean ± SEM; points represent independent donors. Statistical significance was determined by two-tailed t test; *P < 0.05; ns, not significant.

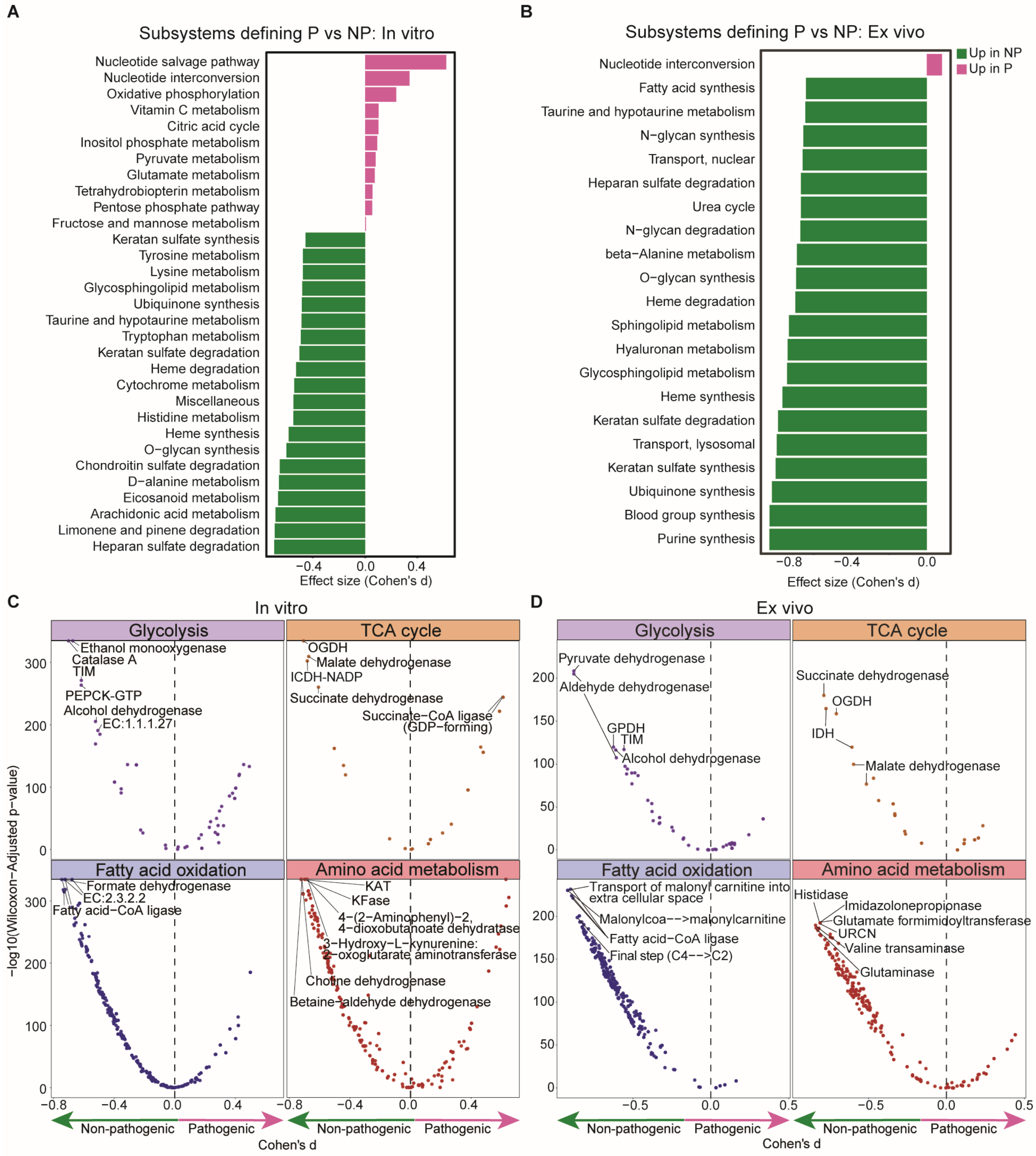


**Supplemental Figure 5. COMPASS identifies metabolic differences between Th17 states in in vitro-skewed and nanovial-enriched samples.** (**A and B**) COMPASS-predicted metabolic subsystems distinguishing Th17p from Th17np cells in *in vitro*-skewed (A) and nanovial-enriched ex vivo (B) samples. Bars show Cohen's d effect sizes. Positive values (magenta) indicate pathways associated with Th17p cells, and negative values (green) indicate pathways associated with Th17np cells. (**C and D**) Reaction-level differences across glycolysis, the TCA cycle, fatty-acid oxidation, and amino-acid metabolism in in vitro-skewed (C) and ex vivo (D) samples. The x axis shows Cohen's d, and the y axis shows −log10 Wilcoxon-adjusted P values. Dashed lines indicate effect-size thresholds; selected reactions are labeled.

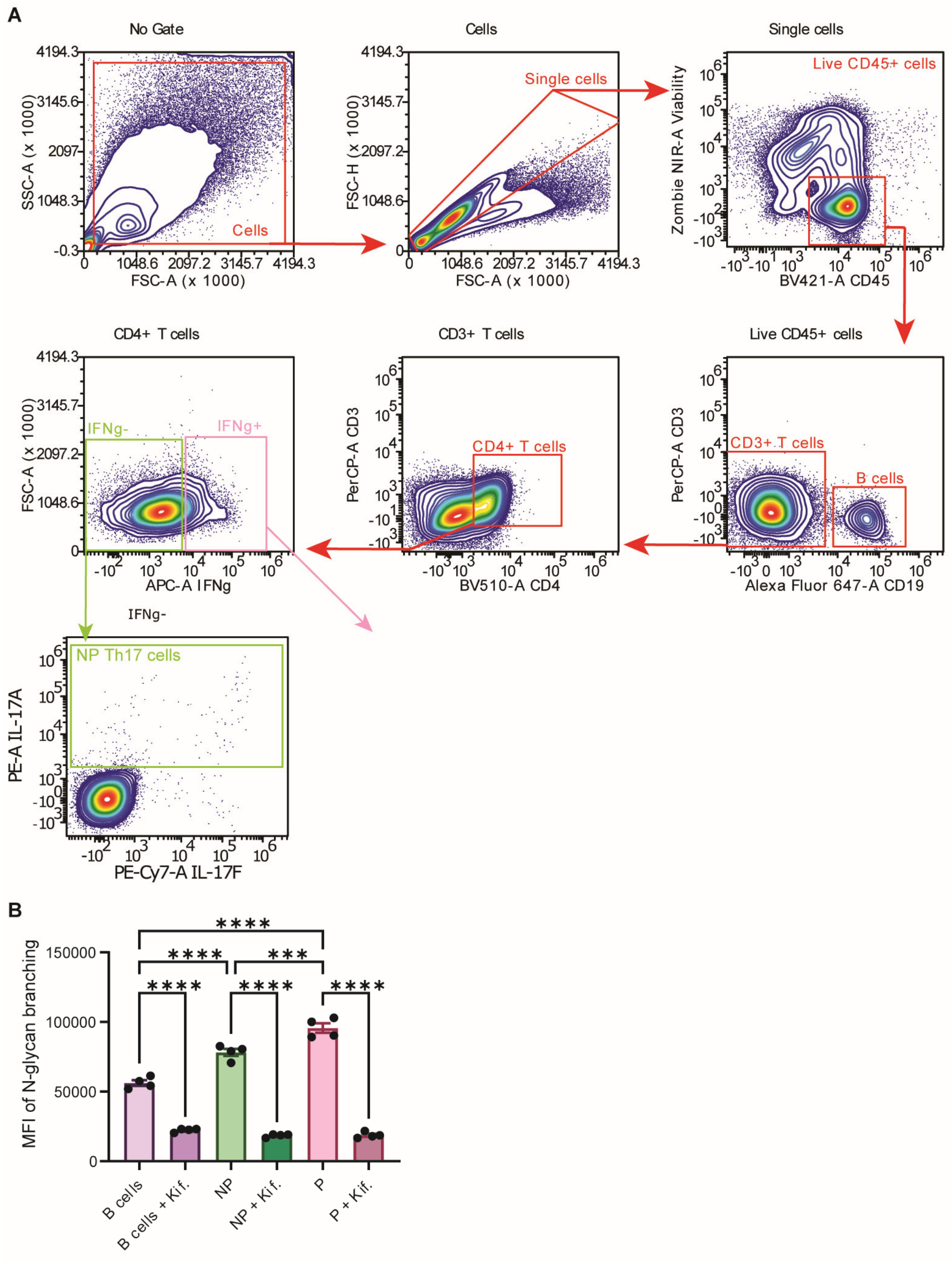


**Supplemental Figure 6. Th17p cells exhibit increased complex N-glycan branching.** (**A**) Representative gating strategy identifying live CD45+ cells, B cells, CD4+ T cells, and Th17np and Th17p subsets for lectin staining. (**B**) Phaseolus vulgaris leukoagglutinin (L-PHA) staining, reported as MFI, in B cells, Th17np cells, and Th17p cells with or without 10 μM kifunensine (Kif), an inhibitor of complex N-glycan formation (n = 4 donors). Bars show mean ± SEM; points represent independent donors. Statistical significance was determined by two-way analysis of variance; ***$P < 0.001$ and ****$P < 0.0001$.

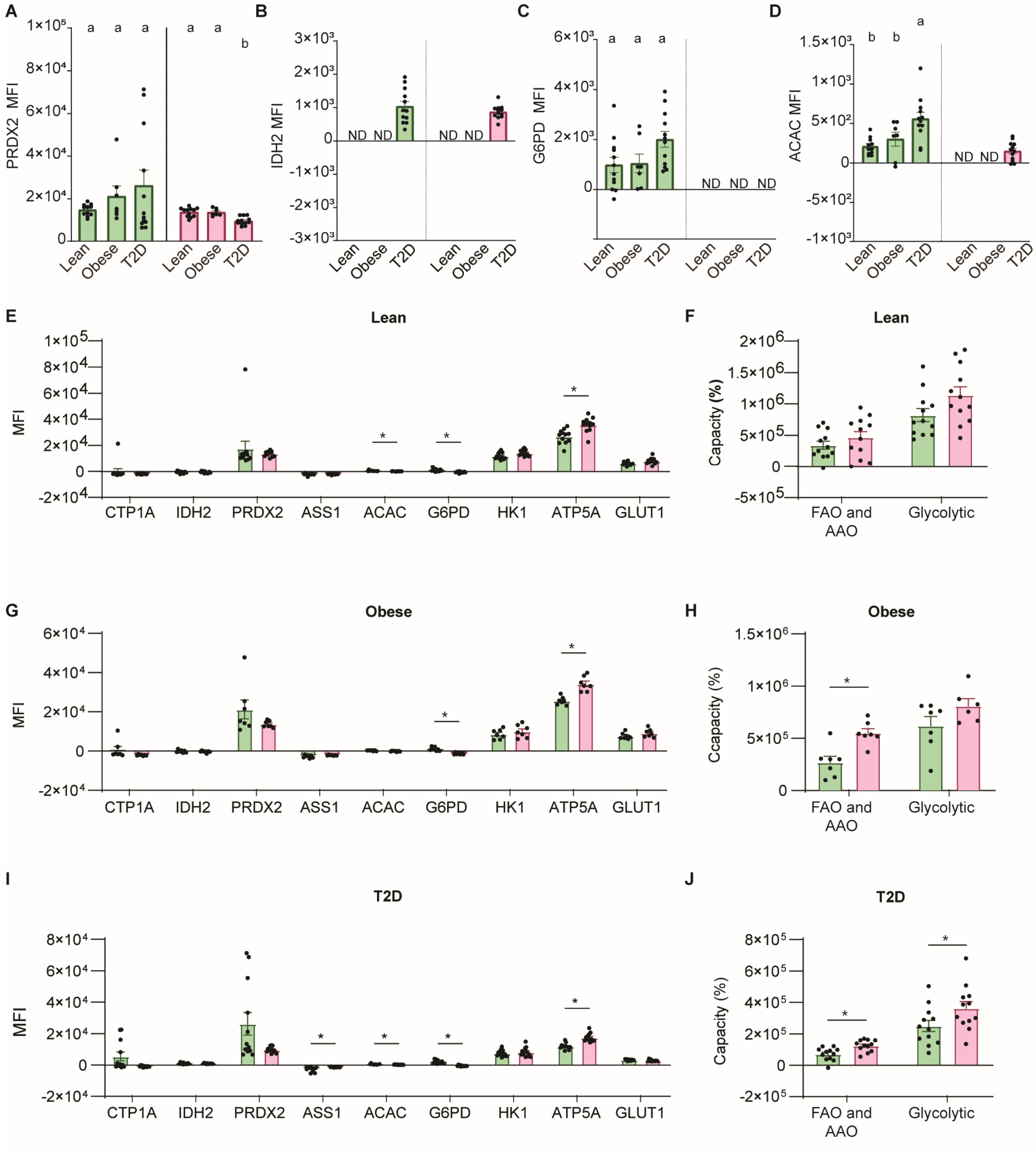


**Supplemental Figure 7. Metabolic features of Th17np and Th17p cells across lean, obese, and type 2 diabetes donors.** (**A to D**) PRDX2 (A), IDH2 (B), G6PD (C), and ACAC (D) MFI in Th17np and Th17p cells from lean, obese, and obese T2D donors. (**E, G, and I**) Comparison of metabolic-protein MFI between Th17np and Th17p cells from lean (E), obese (G), and obese T2D (I) donors. (**F, H, and J**) Comparison of SCENITH-derived FAO and amino acid oxidation (AAO) capacity and glycolytic capacity between Th17np and Th17p cells from lean (F), obese (H), and obese T2D (J) donors. Green and magenta indicate Th17np and Th17p cells, respectively. Bars show mean ± SEM; points represent individual donors. In (A to D), different letters denote statistically significant groups ($P < 0.05$); groups sharing a letter are not significantly different. In (E to J), *$P < 0.05$. ND, not detected.

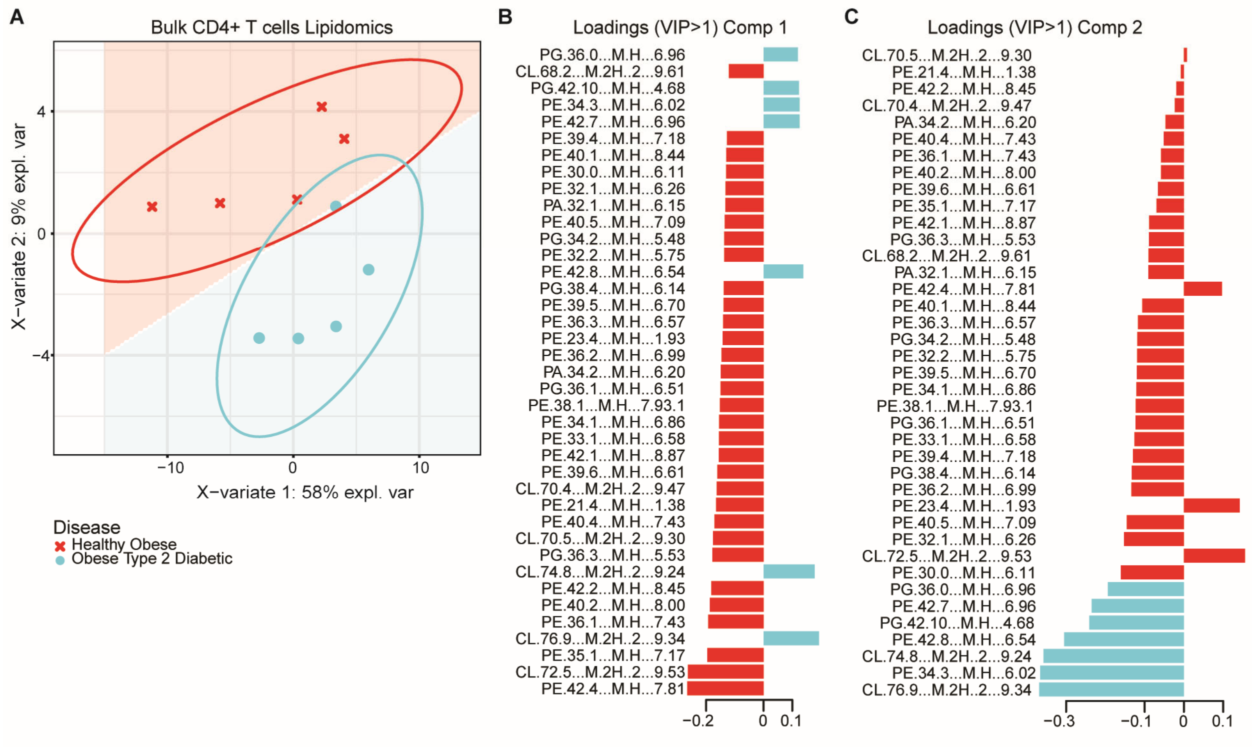


**Supplemental Figure 8. CD4+ T cell lipid profiles differ between obese donors with and without type 2 diabetes.** (**A**) Partial least-squares discriminant analysis (PLS-DA) of untargeted lipidomics from activated bulk CD4+ T cells of normoglycemic obese donors (n = 5) and obese T2D donors (n = 5). Ellipses show confidence regions, and axes indicate the percentage of explained variance. (**B and C**) Variable-importance-in-projection (VIP > 1) loadings for PLS-DA components 1 (B) and 2 (C). Positive and negative loadings indicate the direction of each lipid species' contribution to group separation. Lipid species are labeled by class and detected ion.

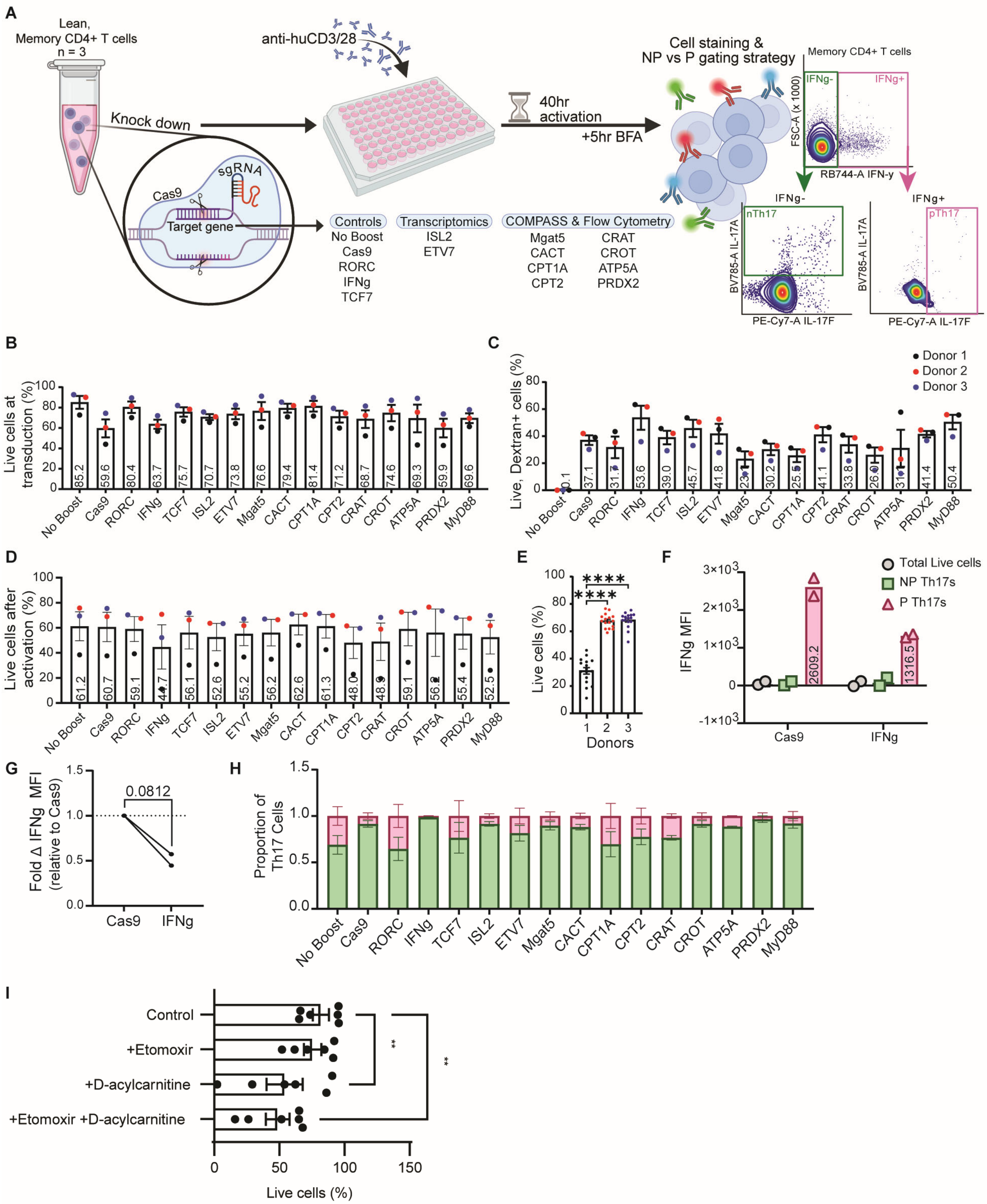


**Supplemental Figure 9. Validation of the targeted CRISPR screen and pharmacologic perturbation of acylcarnitine metabolism.** (**A**) Experimental workflow for CRISPR-Cas9 knockdown of candidate transcriptional and metabolic regulators in memory CD4+ T cells from three healthy lean donors. Ribonucleoprotein complexes were delivered by mechanoporation, followed by 40 hours of activation, brefeldin A during the final 5 hours, and flow cytometry quantification of Th17np and Th17p cells. (**B to D**) Percentage of live cells immediately after mechanoporation (B), live dextran-positive cells as a measure of delivery (C), and live cells after activation (D) for each target. (**E**) Post-activation viability by donor. (**F**) IFN-γ MFI in total live, Th17np, and Th17p cells after Cas9-only control or IFNG knockdown. (**G**) Fold change in total-cell IFN-γ MFI after IFNG knockdown relative to matched Cas9-only controls. (**H**) Proportions of Th17np and Th17p cells among live cells after perturbation of the indicated targets. (**I**) Percentage of live cells after treatment with vehicle, etomoxir, D-acylcarnitine, or etomoxir plus D-acylcarnitine. Bars show mean ± SEM; points represent independent donors. The P value in (G) was determined by paired t test. *P < 0.05, **P < 0.01, and ****P < 0.0001.

**Supplemental Table 1. Clinical demographics of study participants with MS**

| **Characteristic** | **MS**<br>N = 8[1] |
|---|---|
| **Age** | 49 (31 - 57) |
| **Gender** | |
| Female | 6 / 8 (75%) |
| Male | 2 / 8 (25%) |
| **Ethnicity** | |
| | 8 / 8 (100%) |
| **BMI** | 29 (21 - 40) |

[1] Median (range) or Frequency (%)

**Supplemental Table 2. Clinical demographics of study participants stratified by disease status**

| Characteristic | Overall<br>N = 31[1] | Lean<br>N = 12[1] | Obese<br>N = 7[1] | Obese T2D<br>N = 12[1] | p-value[2] |
|---|---|---|---|---|---|
| **Age** | 44 (21 - 67) | 35 (21 - 51) | 38 (27 - 61) | 59 (31 - 67) | 0.002 |
| **Gender** | | | | | 0.16 |
| Female | 12 / 31 (39%) | 3 / 12 (25%) | 5 / 7 (71%) | 4 / 12 (33%) | |
| Male | 19 / 31 (61%) | 9 / 12 (75%) | 2 / 7 (29%) | 8 / 12 (67%) | |
| **Ethnicity** | | | | | 0.012 |
| African American | 5 / 31 (16%) | 4 / 12 (33%) | 0 / 7 (0%) | 1 / 12 (8.3%) | |
| African American Asian/Pacific Islander | 1 / 31 (3.2%) | 1 / 12 (8.3%) | 0 / 7 (0%) | 0 / 12 (0%) | |
| American Indian | 1 / 31 (3.2%) | 0 / 12 (0%) | 1 / 7 (14%) | 0 / 12 (0%) | |
| Asian/Pacific Islander | 7 / 31 (23%) | 3 / 12 (25%) | 0 / 7 (0%) | 4 / 12 (33%) | |
| Hispanic/Latino | 10 / 31 (32%) | 2 / 12 (17%) | 6 / 7 (86%) | 2 / 12 (17%) | |
| Hispanic/Native American | 1 / 31 (3.2%) | 0 / 12 (0%) | 0 / 7 (0%) | 1 / 12 (8.3%) | |
| Other | 1 / 31 (3.2%) | 1 / 12 (8.3%) | 0 / 7 (0%) | 0 / 12 (0%) | |
| White/Non-hispanic | 5 / 31 (16%) | 1 / 12 (8.3%) | 0 / 7 (0%) | 4 / 12 (33%) | |
| **BMI** | 29 (22 - 44) | 24 (22 - 26) | 32 (28 - 38) | 31 (26 - 44) | <0.001 |

[1] Median (range) or Frequency (%)

[2] Kruskal-Wallis rank sum test; Fisher's exact test
